# Corticospinal neuroplasticity and sensorimotor recovery in rats treated by infusion of neurotrophin-3 into disabled forelimb muscles started 24 h after stroke

**DOI:** 10.1101/367573

**Authors:** Denise A. Duricki, Svetlana Drndarski, Michel Bernanos, Tobias Wood, Karen Bosch, Qin Chen, H. David Shine, Camilla Simmons, Steven C.R. Williams, Stephen B. McMahon, David J. Begley, Diana Cash, Lawrence D.F. Moon

**Author notes:** Corresponding author: Dr Lawrence Moon, Neurorestoration Group, Wolfson Centre for Age-Related Diseases, 16 – 18 Newcomen Street, London, SE1 1UL, United Kingdom.

## Abstract

Stroke often leads to arm disability and reduced responsiveness to stimuli on the other side of the body. Neurotrophin-3 (NT3) is made by skeletal muscle during infancy but levels drop postnatally and into adulthood. It is essential for the survival and wiring-up of sensory afferents from muscle. We have previously shown that gene therapy delivery of human NT3 into the affected *triceps brachii* forelimb muscle improves sensorimotor recovery after ischemic stroke in adult and elderly rats. Here, to move this therapy one step nearer to the clinic, we set out to test the hypothesis that intramuscular infusion of NT3 protein could improve sensorimotor recovery after ischemic cortical stroke in adult rats. To simulate a clinically-feasible time-to-treat, twenty-four hours later rats were randomized to receive NT3 or vehicle by infusion into *triceps brachii* for four weeks using implanted minipumps. NT3 increased the accuracy of forelimb placement during walking on a horizontal ladder and increased use of the affected arm for lateral support during rearing. NT3 also reversed sensory deficits on the affected forearm. There was no evidence of forepaw sensitivity to cold stimuli after stroke or NT3 treatment. MRI confirmed that treatment did not induce neuroprotection. Functional MRI during low threshold electrical stimulation of the affected forearm showed an increase in peri-infarct BOLD signal with time in both stroke groups and indicated that neurotrophin-3 did not further increase peri-infarct BOLD signal. Rather, NT3 induced spinal neuroplasticity including sprouting of the spared corticospinal and serotonergic pathways. Neurophysiology showed that NT3 treatment increased functional connectivity between the corticospinal tracts and spinal circuits controlling muscles on the treated side. After intravenous injection, radiolabelled NT3 crossed from bloodstream into the brain and spinal cord in adult mice with or without strokes. Our results show that delayed, peripheral infusion of neurotrophin-3 can improve sensorimotor function after ischemic stroke. Phase I and II clinical trials of NT3 (for constipation and neuropathy) have shown that peripheral, high doses are safe and well tolerated, which paves the way for NT3 as a therapy for stroke.

## Introduction

Ischemic stroke occurs in the brain when blood flow is restricted, causing brain cells to die rapidly. Movements on the opposite side of the body are frequently affected^1^. Stroke victims also often exhibit lack of responsiveness to stimuli on their affected side. The W.H.O. estimates that, worldwide, there are 31 million stroke survivors, with another 9 million new strokes annually. The vast majority of stroke victims are not eligible for the few therapies that improve outcome because they arrive in hospital too late for reperfusion to be effective^2^. Treatment six hours or more after ischemic stroke is usually limited to rehabilitation: therapies that reverse sensory impairments and locomotor disability are urgently needed, and these must work when initiated many hours after stroke.

Neurotrophin-3 is a growth factor which plays a key role in the development, and function of locomotor circuits that express NT3 receptors, including descending serotonergic^3^ and corticospinal tract (CST) axons^4^ and afferents from muscle and skin that mediate proprioception and tactile sensation^5^–^7^. However, peripheral levels of NT3 drop in the postnatal period^8^. We and others had shown that delivery of NT3 into the CNS promotes recovery in rodent models of spinal cord injury^9–18^ but this involved invasive routes of delivery (*e.g*., intraspinal injection or intrathecal infusion) or gene therapy. We also recently showed that injection of an adeno-associated viral vector (AAV) encoding full-length human NT3 (preproNT3, 30kDa) into forelimb muscles 24 hours after stroke in adult or elderly rats improved sensorimotor recovery^19^. We had originally expected that AAV1 would be trafficked from muscle to the spinal cord retrogradely in axons^20^ and that this would enhance secretion of NT3 by motor neurons, leading to sprouting of the spared CST^11,21,22^ and sensorimotor recovery. Although NT3 protein was overexpressed in injected muscles, to our surprise we found little evidence for expression of the human NT3 transgene in the spinal cord or cervical DRGs^17,19^ using this dose and preparation of AAV. This serendipitous result led us to reject our original assumption that sensorimotor recovery required expression of the human NT3 transgene in the spinal cord and to wonder whether peripheral infusion of NT3 protein would suffice.

Accordingly, here, we test the hypothesis that infusion of the mature form of the NT3 protein (14 kDa) into disabled forelimb muscles improves sensorimotor recovery. This is consistent with work by others^23^ including a study showing that a signal from muscle spindles can improve neuroplasticity of descending pathways and can enhance recovery after CNS injury^24^. Notably, NT3 protein is synthesised by muscle spindles^25^ and can be transported from muscle to sensory ganglia and spinal motor neurons in nerves^7^,^17^,^19^ and from the bloodstream to the CNS^26–28^. This route of administration and time frame is clinically feasible so to take this potential therapy one step nearer the clinic, we next set out to determine whether intramuscular infusion of human NT3 protein (mature form, 14kDa) would improve outcome after stroke (*i.e*., bypassing the use of gene therapy and spinal surgery).

Importantly, the mature form of the NT3 protein has excellent translational potential: Phase I and II clinical trials have shown that repeated, systemic, high doses of NT3 protein are well-tolerated, safe and effective in more than 200 humans with sensory and motor neuropathy (Charcot-Marie-Tooth Type 1A)^29^ or constipation including in people with spinal cord injury^29–33^. In contrast to other neurotrophins, NT3 does not cause any serious adverse effects such as pain^34^ probably because its principal high affinity receptor TrkC is not expressed on adult nociceptors^6^,35. These studies pave the way for NT3 as a therapy for stroke in humans. We now show in a blinded, randomized preclinical trial that treatment of disabled upper arm muscles with human NT3 protein reverses sensory and motor disability in rats when treatment is initiated in a clinically-feasible timeframe (24 hours after stroke).

## Methods

### Experimental design

Rats received unilateral focal cortical stroke or underwent sham surgery^19,36^ (Fig. 1a,b). Twenty-four hours after stroke, rats were allocated to treatment using NT3 or vehicle, infused into affected *triceps brachii* muscles for one month *via* implanted catheters and subcutaneous osmotic minipumps. Experiments were performed in accordance with guidelines from the Stroke Therapy Academic Industry Roundtable (STAIR) and others and our findings were reported in accordance with the ARRIVE (Animals in Research: Reporting *In Vivo* Experiments) guidelines. All surgical procedures, behavioural testing and analysis were performed using a randomised block design. All surgeries, behavioural testing and analysis were performed with investigators blinded to treatment groups. Rats were randomised to surgery by drawing a rat identity number from an envelope and then a stroke/sham allocation from an envelope. Allocation concealment was performed by having NT3 and vehicle stocks coded by an independent person prior to loading pumps. Behavioural testing was conducted blind and codes were only broken after behavioural analyses were complete. 65 Lister hooded (∼4 months; 200–300 g) outbred female rats (Charles River, UK) and 95 adult C57Bl/6 mice (7–8 weeks) were used. All procedures were in accordance with the UK Home Office guidelines and Animals (Scientific Procedures) Act of 1986.

**Figure 1:**
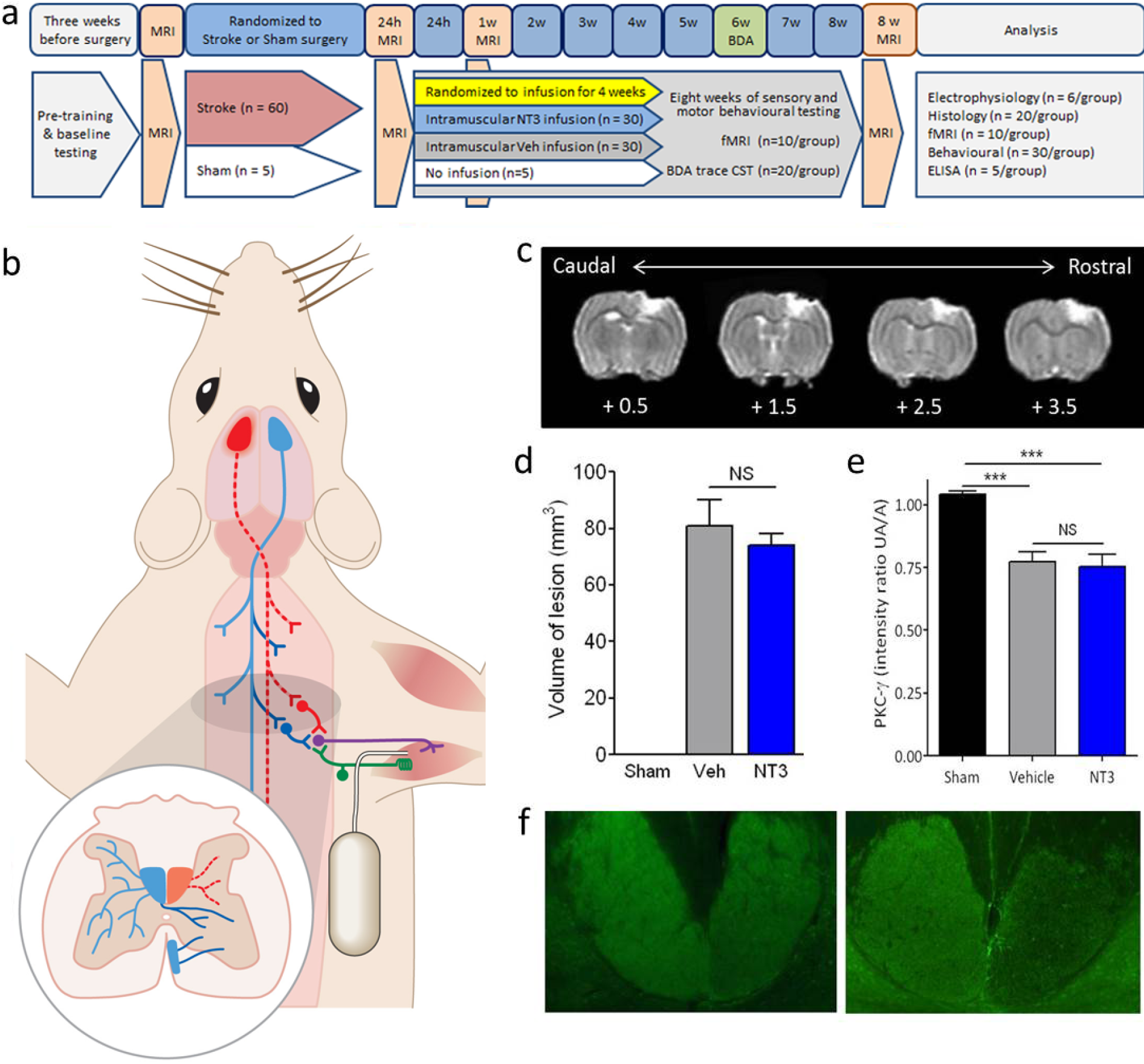
Focal stroke caused unilateral infarction in sensorimotor cortex, resulting in a loss of corticospinal axons. **a, b)** Rats received sham surgery or unilateral cortical stroke (red) and 24 hours later infusion of NT3 or vehicle into the disabled *triceps brachii* was initiated for one month. Six weeks after stroke, anterograde tracer was injected into the contralesional hemisphere (blue). Rats underwent 8 weeks of behavioural testing. Structural MRI was conducted on all rats at 24 hours, 1 week and 8 weeks after stroke and fMRI was conducted in a subset of rats at baseline, week 1 and week 8. Electrophysiology was performed in the subset of rats which did not receive BDA tracing. All surgeries, treatments and behavioural testing were performed using a randomized block design and the study was completed blinded to treatment allocation. **c)** T2 MRI scans 24 hours after stroke, immediately prior to treatment, showing infarct in coronal sections rostral (mm) to bregma. **d)** There were no differences between stroke groups in mean lesion volume at 24 hours (Mann Whitney p values = 0.86). **e)** Photomicrographs showing loss of CST axons in the upper cervical dorsal columns 8 weeks after stroke (right) relative to sham surgery (left), visualised using PKCγ immunofluorescence. **f)** Stroke caused a significant loss of CST axons relative to shams in the dorsal columns (Kruskal Wallis χ^2^=12.0, p=0.002; Mann Whitney p-values<0.001) with no differences between vehicle and NT3 treated rats (Mann Whitney p=0.96). Image shows one representative stroke animal and one representative sham animal.

### Stroke or sham lesions

Rats were maintained (Specific Pathogen Free) in groups of 2 to 4 in Plexiglas housing with tunnels and bedding on a 12:12 hour light/dark cycle with food and water *ad libitum*. Focal ischemic stroke was induced in the hemisphere representing the dominant forelimb (Supplementary Fig. 1), as determined by the cylinder behavioural test. Stroke lesions (*n* = 60) were performed as previously described^19^. Briefly, animals were transferred to a stereotaxic frame (David Kopf Instruments, USA) where a midline incision was made, the cortex was then exposed via craniotomy using the following co-ordinates [defined as anterioposterior (AP), mediolateral (ML)]: AP 4 mm to – 2 mm, ML 2 mm to 4 mm, relative to bregma. Endothelin-1 (ET-1, 400 pmol/µL in sterile saline; CalBioChem) was applied using a glass micropipette attached to a Hamilton syringe. 1 μl of ET-1 to was applied to the overlying dura to reduce bleeding, and immediately thereafter, the dura mater was incised and reflected. Four 1 μl volumes of ET-1 were administered topically and four 1 μl volumes were microinjected intracortically (at a depth of 1 mm from the brain surface) at the following co-ordinates (from bregma and midline, respectively):

AP + 3.5 mm, ML 2.8 mm

AP + 2.0 mm, ML 2.8 mm

AP + 0.5 mm, ML 2.8 mm

AP – 1.0 mm, ML 2.8 mm

Temperature was maintained using a rectal probe connected to a homeothermic blanket (Harvard Apparatus, USA) placed under the animal which maintained rectal temperature at 36 ± 1°C. Prior to suturing, the animal was left undisturbed for 5 minutes. Modifying previous work^36^, the skull fragment was then replaced and sealed using bone wax (Covidien, UK). 100% (60/60) of rats survived this stroke surgery. Sham-operated (*n*= 5) rats received all procedures up to, but not including, craniotomy or endothelin-1 injection. Animals were given buprenorphine (0.01 mg/kg, subcutaneously) for postoperative pain relief.

Our method of inducing stroke with ET-1 is advantageous for evaluating regenerative stroke therapies for four reasons: 1) our model produces ischemic lesions that model small focal human strokes rather than larger “malignant” strokes that tend to be fatal in humans^37^; 2) our model targets sensorimotor cortex; 3) our stroke model involves only low mortality rates and has reasonable reproducibility^36^ with a proven ability to detect therapies that induce neuroplasticity and functional recovery^19^,38; 4) our model causes sustained sensorimotor deficits (*e.g*., impaired use of limbs) which are common neurological symptoms of human stroke.

### Magnetic resonance imaging

Structural images were obtained prior to stroke, 24 hours after stroke and at one and eight weeks after stroke. MR imaging was conducted on a 7 Tesla (T) horizontal bore VMRIS scanner (Varian, Palo Alto, CA, USA). Animals were anaesthetised using 3.5% isoflurane, in 0.8 L/min medical air and 0.2 L/min medical oxygen in an induction chamber. Once anaesthetised they were secured in a stereotaxic head frame inside the quadrature birdcage MR coil (43 mm internal diameter, ID) and placed into the scanner. Each animal’s physiology was supervised throughout the procedure using a respiration monitor (BIOPAC, USA) and a pulse oximetry sensor (Nonin, USA) that interfaced with a PC running Biopac. Additionally, an MRI compatible homeothermic blanket (Harvard Apparatus, USA) placed over the animal responded to any alterations in the body temperature identified by the rectal probe, and maintained temperature at 37 ± 1°C. The T2 weighted MR images were acquired using a fast spin-echo sequence: effective echo time (TE) 60 ms, repetition time (TR) 4000 ms, field of view (FOV) 40 mm × 40 mm and an acquisition matrix 128 × 128, acquiring 20 × 1 mm thick slices in approximately 8 minutes. At the end of the study (to avoid affecting blinding or randomization), lesion volumes at the 24 hour time point were measured using a semi-automatic contour method in Jim software under blinded conditions (Xinapse systems Ltd).

### Functional magnetic resonance imaging (fMRI)

Functional magnetic resonance imaging (fMRI) was performed in a subset of rats that did not receive intracortical injections of BDA tracer (n = 10/group). Images were acquired prior to stroke and at one and eight weeks after stroke during non-noxious somatosensory stimulation of the affected or less-affected wrist^19^. This involves delivery of small electrical currents to a wrist whilst the subjects were kept anaesthetised using medium dose Alpha-Chloralose suitable for recovery and longitudinal (repeated) imaging^39^. Alpha chloralose anesthesia was prepared by mixing equal amounts of Borax Decahydrate (Sigma) and alpha chloralose-Pestanal (analytical standard, <8% beta isoform, 45373, Sigma, UK) in physiological saline each at a concentration of 40 mg/ml in a glass beaker at 52°C prior to filtering using a 0.22 μm filter.

Rats were first anaesthetised using 4% isoflurane, 0.8 L/min medical air and 0.2 L/min oxygen. A tail cannulation was performed and the animal was transferred to the MRI machine. A bolus of 40 mg/kg alpha chloralose-Pestanal was injected intravenously and then the isoflurane was switched off after 5 minutes. An infusion line for continuous application of alpha chloralose was then attached to the cannula at 10 mg/kg/h over the experimental time. Medical air (0.8 L/min) and oxygen (0.2 L/min) was continuously delivered throughout the scanning period. MR images were obtained using a 7 Tesla scanner (Varian, Agilent). Initially, a T2-weighted structural scan was acquired, using a fast spin echo (FSE) sequence with repetition time (TR) = 4000 ms, echo time (TE) = 60 ms and field of view (FOV) of 40 × 40 mm, yielding 20 slices with voxel size of 0.3215 × 0.3125 × 1 mm in approximately 8 min. fMRI scans were acquired using a Multi-Gradient-Multi-Echo sequence (TR = 360 ms, TEs = 5, 10, 15 ms, voxel size 0.5 mm × 0.5 mm × 1 mm, resolution 64 × 64 × 20, scan time 23 s). 100 volumes were acquired with a pseudo-random onoff stimulation of the forepaw at 3 Hz (400 μs, 2 mA pulse) using a platinum subdermal needle electrode (F-E2, Grass Technologies, USA) and a TENS (Transcutaneous Electrical Nerve Stimulation) pad. It has previously been shown that the use of a TENS pad results in this intensity of stimulation being innocuous rather than noxious: whereas blood pressure is not altered at 2 mA stimulation, it is increased with 3.8 mA stimulation (see references in^19^). The order of paw stimulation was also randomised. Animals were closely monitored following the end of the scanning, given 5 ml of saline (*subcutaneous*, room temperature) and were kept in a warmed incubator (31 deg C) until fully conscious: this takes 2 to 3 hours due to slow pharmacokinetics of alpha chloralose. Two animals died following alpha chloralose anaesthesia due to breathing difficulties in recovery. Scans with obvious imaging artefacts were discarded, leaving final group numbers of n=7, 9, 7 and n=9, 8, 4 at weeks 0, 1 and 8 for NT3 and vehicle treated groups respectively. The resulting images were analyzed with SPM-8 (Statistical Parametric Mapping, FIL, UCL). In order to make sure all lesions were in the same side of the brain, images with right-hand side strokes were rotated about the sagittal mid-plane, so that the lesioned hemisphere always appeared on the left. Functional scans were initially realigned to the first image in the time-series in order to correct for movements of the head. The first volume of the functional scan was then spatially registered to the structural image, which was, in turn, linearly warped to a template brain. Linear warping was used in this step in order to avoid deforming the lesion region. Warping parameters obtained during registration of structural image to template were applied to the realigned functional time-series, resulting in structural and functional images that are all in a standard space. Finally, functional images were smoothed using a Gaussian kernel with Full Width Half Maximum of 1.0 × 1.0 × 2 mm (twice the voxel size). Because of the relatively long effective TR of the functional images, a PET basic model (one-sample t-test) was used for first-level analyses with covariates consisting of the pseudo-random stimulation pattern (paradigm), and the estimated movement parameters of each individual rat. Volumes signal intensity was globally scaled and individual masks, generated from the fast spin echo (FSE) structural scan for each rat brain at each time-point using a 3D Pulse-Coupled Neural Network^40^ (also registered to the template), were used as explicit masks for the first-level statistical analysis. Contrast images from the First-level analysis were then carried onto a Second-level (random effects) group analysis. Effects of group (i.e. NT3 or vehicle treated) and stimulated paw (i.e. affected or less-affected) were used to create statistical comparisons. A flexible factorial analysis was used to compare the difference between the NT3 and vehicle treated groups in the change from baseline to 8 weeks^41^. Statistical parametric maps were generated using an uncorrected threshold of p < 0.05; images show group mean activations and T values are given.

### NT3 or vehicle treatment

24 hours after stroke (immediately after MRI), rats were allocated to treatment using a randomized block design. Allocation concealment was performed by having NT3 and vehicle stocks coded by a third party. Rats were anaesthetised as above and a small incision was made between elbow and axilla, and a small subcutaneous space was formed to the lower back. The osmotic pump with the catheter attached was positioned in this subcutaneous space and then an ultrafine, flexible catheter was implanted approximately 15 millimetres into the proximal end of the long head of the *triceps brachii* muscle on the disabled side and this was sutured in place (Prolene 5/0, Ethicon, UK). *Triceps brachii* was selected as the site for infusion because this large muscle is involved in forelimb extension during walking and for postural support during rearing) rearing. Note that in *the triceps brachii*, the end plates are located in the belly of the muscle^42^. The catheter was made from stretchable and flexible silicone tubing (ID = 0.012 inches, wall thickness = 0.007 inches, Trelleberg, SF Medical, UK, SFM1–1050) attached to the osmotic pump *via* larger glued-on tubing (ID 0.76 mm, VWR International). A second section of this stiff tubing (10 millimetres long) was inserted to guide the flexible catheter into the triceps; the guide was slid back after the silicone tube was implanted. Catheters were connected to subcutaneous osmotic minipumps (2ML2, Alzet) containing either vehicle (0.9% saline containing 0.1% bovine serum albumin; Sigma; A3059; 9048–46-8) or vehicle containing recombinant human NT3 (kind gift of Genentech Inc., USA). Pump flow moderators were MR-compatible (PEEK micro medical tubing, Durect, #0002511). The original vials contained 0.586 mg/ml recombinant human (“rhu”) NT3 in 20 mM acetate, 140 mM NaCl, pH 5.0. SDS PAGE and proteomic analysis indicates that this is the 119 amino acid mature NT3 protein obtained after proteolytic cleavage of the 258 amino acid proneurotrophin-3 (Uniprot P20873) (Supplementary Figures 7 and 8). The NT3 dose (12 μg/24 hours) was selected based on previous experiments^9^. Pumps were replaced after two weeks and removed after four weeks. Skin was sutured and analgesic administered as above. All rats survived this surgery. Sham rats did not undergo this surgery. Pilot experiments showed that the pump flow rate (5 μl/hour) was sufficient to deliver substances (0.9% saline containing 1% Fast Green, Sigma) to the entire volume of the triceps muscles.

### Enzyme Linked Immunosorbent Assay (ELISAs)

Rats (n = 5/group) were terminally anesthetised (4 weeks following stroke, before pumps were removed) and *triceps brachii* and C7 spinal cord were rapidly dissected and snap frozen in liquid nitrogen prior to storage at –80°C. Tissue was homogenised in ice cold lysis buffer containing 137 mM NaCl, 20 mM Tris-HCl (pH 8.0), 1% NP40, 10% glycerol, 1 mM phenylmethanesulfonylfluoride, 10 μg/ml aprotinin, 1 μg/ml leupeptin and 0.5 mM sodium vanadate, using approximately 10 times the volume of buffer to the wet weight of tissue (10 μl/mg tissue). Protein content was measured and NT3 ELISA was carried out according to manufacturer’s instructions (Emax, Promega).

*n.b*., Promega NT3 ELISA kits are no longer available. However, it was recently discovered that when performing ELISA using other NT3 kits (R&D using “Reagent Diluent” and Abcam using diluent A), measurements of NT3 from skeletal muscle lysates do not provide reliable quantitative data. This is due to so-called “matrix effects” as shown by poor recovery of spiked-in NT3 (<80% or >120%) and non-linear relationship between concentration of input material and estimated NT3 concentration based on a dilution series of muscle homogenate. To overcome the effect of interfering substances, samples should be diluted and appropriate diluents to prepare standards and diluted samples must be used. The Abcam kit used with diluent B (not A), however, provided reliable quantitative results (Dr. Aline Barroso Spejo, unpublished results).

### Behavioural testing

We assessed sensory and motor deficits after stroke using the cylinder test (to assess postural support by forelimbs during rearing), adhesive patch test (to assess responsiveness to tactile stimuli), horizontal ladder (to assess forelimb and hindlimb skilled locomotion), a grip strength test and a test used to monitor unusual responses to cold stimuli (cold allodynia)^19^,38. All behavioural testing was carried out by an experimenter blinded to surgery and treatment groups. Rats were handled and trained for three weeks on the horizontal ladder before the study began. Preoperative baseline scores for the horizontal ladder, the vertical cylinder and the grip strength test were collected one week before surgery.

#### Assessment of somatosensory responsiveness

The “adhesive patch” test was used to measure 1) the time taken to contact stimuli on the wrists, 2) the time taken to remove stimuli from the wrists, and 3) the magnitude of lack of responsiveness to stimuli on the affected wrist^32,34,46,47^. For each trial, a round adhesive patch (13 mm diameter, Ryman) was applied to each wrist on the dorsal side and the animal was returned to its home cage. Two times were recorded for both forepaws: (1) contact and (2) remove; where “contact” represents the time taken for the animal to contact an adhesive patch with its mouth, and “remove” represents the time taken for the animal to remove the first adhesive patch from its wrist. To determine whether the rats preferentially removed a sticker from their less-affected wrist before their more-affected wrist, the order and side of label removal was recorded. This was repeated four times per session until a >75% preference had been found; if this was not the case a fifth trial was conducted. The magnitude of asymmetry was established using the seven levels of stimulus pairs on both wrists as previously described (Figure 2a). From trial to trial, the size of the stimulus was progressively increased on the affected wrist and decreased on the less affected wrist by an equal amount (14.1 mm^2^), until the rat removed the stimulus on the affected wrist first (reversal of original bias). The higher the score, the greater the degree of somatosensory impairment.

**Figure 2:**
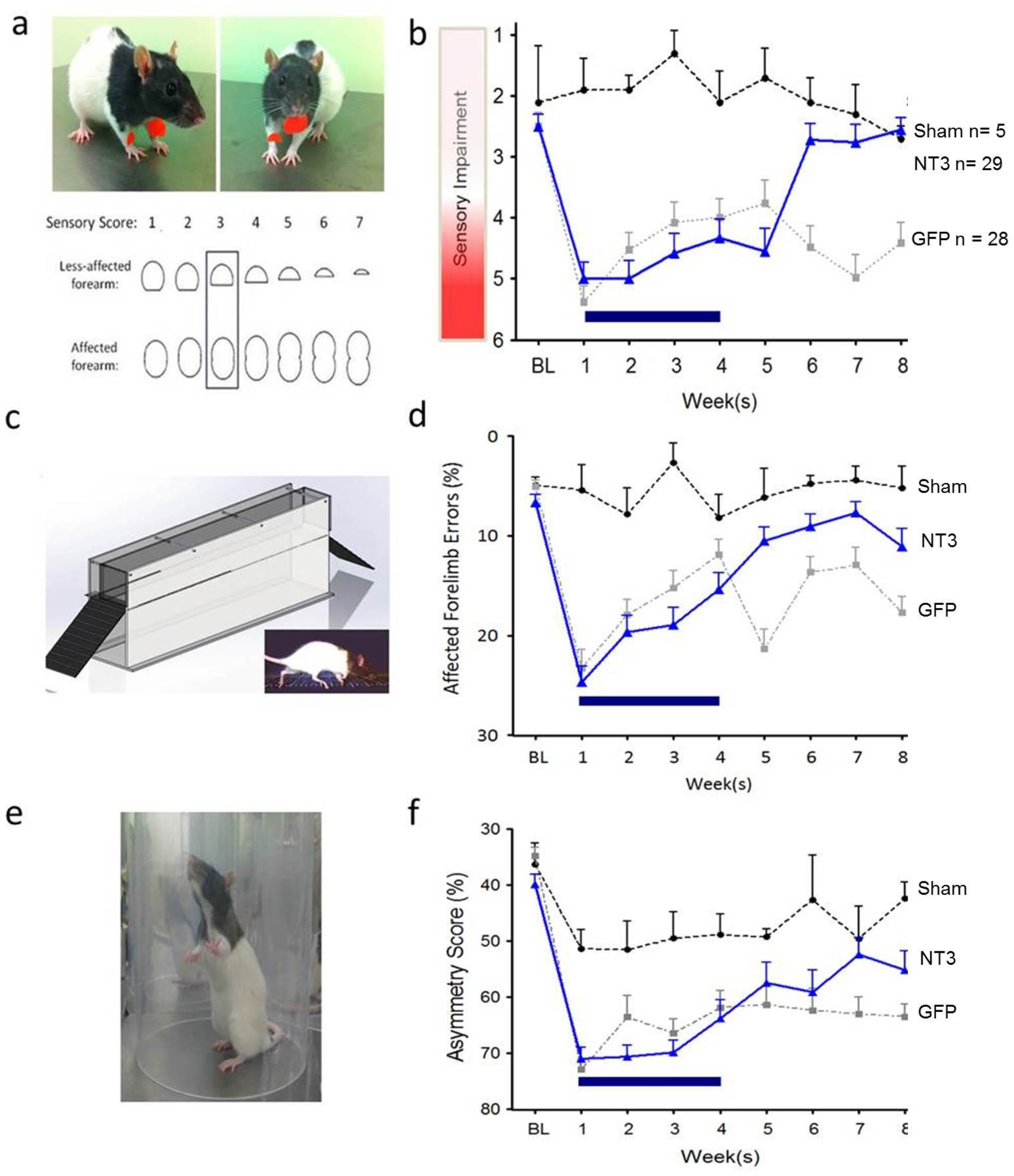
Delayed NT3 treatment improved responsiveness to somatosensory stimulation, improved walking and partially restored use of the affected forelimb for lateral support during rearing. **a-c)** Somatosensory deficits were assessed using pairs of adhesive patches attached to the rat’s wrists. **b)**Treatment with NT3 caused improvement compared to vehicle (linear model; F2,126=32.5, p<0.001; *post hoc* p=0.003): whereas vehicle-treated stroke rats showed a deficit relative to sham rats which persisted for eight weeks (linear model; F14,263=4.1, p<0.001; *post hoc* p values<0.02), NT3 caused improvements such that by six weeks they had recovered fully relative to shams (*post hoc* p values>0.4) and to vehicle-treated stroke rats (*post hoc* p values<0.001). There were no differences between NT3 and vehicle treated rats at one week (t-test p=0.33). **c)** Accuracy of paw placement by the affected forelimb during walking was assessed using a horizontal ladder with irregularly spaced runs. **d)** One week after stroke, NT3 and vehicle treated rats made a similar number of misplaced steps (t-test p=0.78), expressed as a percentage of total steps. Importantly, the NT3 group progressively recovered compared to the vehicle group (group F_2,52_=17.0, p<0.001; *post hoc* p=0.008) and differed from the vehicle group from weeks 5 to 8 (group × time F14,355=6.2, p<0.001; *post hoc* p values<0.005) and whereas the vehicle group remained impaired relative to shams from weeks 5 to 8 (p values<0.006), from weeks 5 to 8 the NT3 group made no more errors than shams (*post hoc* p-values>0.05). **e)** The vertical cylinder test assessed use of the affected forelimb for lateral support during rearing. **f)** Stroke caused a reduction in the use of the affected forelimb during rearing in a vertical cylinder in both NT3 and vehicle treated rats relative to shams (group F_2,56_=6.1, p=0.004; *post hoc* p values=0.001 and 0.003, respectively) with no differences between stroke groups at one week (p=0.52). NT3 treatment caused a progressive recovery in the use of the affected forelimb (group × wave F14,356=1.78, p=0.04) by weeks 7 and 8 relative to vehicle treated rats (p values = 0.015 and 0.067, respectively). Sham group n=5, Stroke and NT3 group n=29, Stroke and vehicle group=28.

#### Walking

To assess impairments in forelimb and hindlimb function after stroke, rats were videotaped as they walked along a horizontal ladder. Rats were videotaped crossing a horizontal ladder (1 m) with irregularly spaced rungs (1 to 4 cm spacing changed weekly) weekly, 3 times per session. Any slight paw slips, deep paw slips and complete misses were scored as errors.

The mean number of errors per step was calculated for each limb for each week (Foot faults are routinely normalized “per step” after stroke although analysis of foot fault data with or without normalization led to the same conclusions being drawn).

#### Assessment of forelimb use during rearing

The cylinder test was used to assess asymmetries in forelimb use for postural support during rearing within a transparent 20 cm diameter and 30 cm high cylinder^19^. An angled mirror was placed behind the cylinder to allow movements to be recorded when the animal turned away from the camera. During exploration, rats rear against the vertical surface of the cylinder. The first forelimb to touch the wall was scored as an independent placement for that forelimb. Subsequent placement of the other forelimb against the wall to maintain balance was scored as “both.” If both forelimbs were simultaneously placed against the wall during rearing this was scored as “both.” A lateral movement along the wall using both forelimbs alternately was also scored as “both.” Scores were obtained from a total number of 10 full rears to control for differences in rearing between animals. Once scores had been acquired, forelimb asymmetry was calculated using the formula: 100 × (ipsilateral forelimb use + 1/2 bilateral forelimb use)/total forelimb use observations (Hsu and Jones 2005).

#### Grip strength

Forelimb strength was measure using a grip strength meter (Supplementary Figure 3a; MJS Technology, UK). Both the affected and less affected forelimb strength were each measured (simultaneously) at baseline, week 1, week 4 and week 8 following stroke. A pair of force transducers were used in parallel to measure the peak force achieved by a rat’s forelimbs as its bilateral grip was broken by the experimenter gently pulling the rat by the base of the tail horizontally away from the transducer. The average of three strength readings was noted down per session and an average taken for both arms. The difference in grip strength was taken by subtracting the affected forepaw grip strength from the less affected grip strength.

#### Assessment of cold allodynia

The presence or absence of cold allodynia was assessed using standard methods^43^. Rats were placed in a transparent cylinder (20 cm diameter, 30 cm height) atop a mesh wire floor. A drop (50 μl) of acetone was placed against the centre of the forepaw. In the following 20 s after acetone application the rat’s response was monitored. If the rat did not withdraw, flick or lick its paw within this 20 s period then no response was recorded for that trial (0, see below). However, if within this 20 s period the animal responded to the cooling effect of the acetone, then the animal’s response was assessed for an additional 20 s, a total of 40 s from initial application. The reasons for taking a longer period of time to assess the evoked behaviour were to measure only pain-related behaviour evoked by cooling and not startle responses that can occur following the initial application of acetone^43^. Moreover, the behaviour evoked by acetone is often an interrupted series of behaviours, thus it is important to give enough time to see all pain-related behaviours. Responses to acetone were graded to the following 4-point scale: 0, no response; 1, quick withdrawal, flick or stamp of the paw; 2, prolonged withdrawal or repeated flicking of the paw; 3, repeated flicking of the paw with licking directed at the ventral side of the paw. Acetone was applied alternately three times to each paw and the responses scored categorically. Cumulative scores were then generated by adding the scores for each rat together, the minimum score being 0 (no response to any of the six trials) and the maximum possible score being 9 per forepaw.

### Anatomical tracing

To visualize uninjured CST axons, six weeks after stroke, 10% biotinylated dextran amine (BDA; 10,000 MW, Invitrogen) in PBS (pH 7.4) was microinjected unilaterally into the uninjured sensorimotor cortex. Animals were placed in a stereotaxic frame and six burr holes were made into the skull at the following coordinates (defined as anterioposterior (AP), mediolateral (ML): 1) AP: +1 mm, ML: 1.5 mm; 2) AP: +0.5 mm, ML: 2.5 mm; 3) AP: +1.5 mm, ML: 2.5 mm; 4) AP: +0.5 mm, ML: 3.5 mm; 5) AP: +2.0 mm, ML: 3.5 mm; 6) AP: – 0.5 mm, ML: 3.5 mm, relative to bregma. At each site, 0.5 μl injections of BDA (10% in PBS) were delivered using a glass micropipette attached to a Hamilton syringe inserted 2 mm from the skull surface and delivered at a rate of 0.25μl/min. Animals were subsequently left for 2 weeks before being perfused. Tract tracing was not performed in rats that were to undergo functional MRI or neurophysiology.

### Neurophysiology

As described below, we recorded from the ulnar nerve on the affected side and stimulated the ipsilateral median nerve^44^ or, in the pyramids, the corticospinal tract corresponding to the affected or less-affected hemisphere. At the end of the study, 18 rats (6 rats per group) were anaesthetised with an intraperitoneal injection of 1.25g/kg urethane (Sigma-Aldrich). The rat was kept at 37°C with a homeothermic blanket system and rectal thermometer probe. Tracheotomy was performed and a tracheal cannula inserted. The pyramids were then exposed ventrally by blunt dissection and removal of a small area of bone. The brachial plexus of the affected forelimb was exposed from a ventral approach by dissecting the *pectoralis major*. The ulnar and median nerves were dissected free from surrounding connective tissue and cut distally (to prevent twitches of target muscles). Skin flaps from the incision formed a pool, which was filled with paraffin oil.

#### Spinal reflexes

The median and ulnar nerves supply flexor muscles in the forearm, wrist and hand. Stimulation of afferents in the median nerve can generate responses in the ulnar nerve motor neurons. The proximal segment of each nerve was mounted on a pair of silver wire hook electrodes (with >1 cm separation). Electrical stimuli of increasing amplitude from 50 μA to 500 μA, in 50 μA steps, (single 100 μs square wave pulse at 0.5 Hz) were delivered from a constant current stimulator (NL800A Neurolog, Digitimer) to the proximal segment of the cut median nerve. Ulnar nerve responses to each stimulus were recorded from the pair of silver wire hook electrodes connected to a differential pre-amplifier and amplifier (Digitimer) coupled via a PowerLab (AD Instruments) interface to a personal computer running LabChart and Scope software (AD Instruments). An average of 64 sweeps at 400 μA was calculated online for each nerve and used to find the difference in amplitudes of monosynaptic reflexes evoked by median nerve stimulation.

This was achieved using software to calculate the absolute integral of any response between 1.5 ms and 3 ms, regardless of whether a response was observed qualitatively.

#### CST stimulation experiment with ulnar recordings

Ulnar nerve recordings were obtained during stimulation of each pyramid in turn. The concentric bipolar stimulation electrode (FHC CBBPC75) was located 1 mm lateral to the midline and gently lowered through the pyramid up to a maximum depth of 1.5 mm while stimulating at 300 μA (4 pulses, pulse width 100 μs; frequency 500 Hz). At the electrode location providing maximal ulnar nerve response, stimuli of increasing intensity were applied in the range of 50 μA to 400 μA, in 50 μA increments. Five sweeps were captured at each stimulus intensity. The number of spikes 50% greater than the noise, and falling between 17 and 45 ms, was calculated for each sweep. The average number of spikes for 5 sweeps at each amplitude was calculated and the difference in the number of spikes elicited by stimulation of the pyramids from the lesioned or contralesional hemisphere. Furthermore, the signal was rectified and the area under the curve was measured between 17 and 45 ms for each sweep and averaged for the 5 sweeps at each intensity. Each parameter was analysed using twoway repeated measures ANOVA. Graphs show mean and standard error of the mean for the area under the curve for stimuli given at 400 μA.

### Histology

Eight weeks after stroke surgery and two weeks after injection of BDA, rats were terminally anesthetized with sodium pentobarbital (80 mg/kg; i.p.) and perfused transcardially with PBS for 5 minutes, followed by 500 ml of 4% paraformaldehyde in PBS for 15 minutes. The brain, C6-C8 spinal cord, C7 and C8 DRGs and both arms were carefully dissected and stored in 4% paraformaldehyde in PBS for 2 hours and then transferred to 30% sucrose in PBS and stored at 2–5°C. Spinal cord segments C1 and C7 was embedded in OCT and 40 μm transverse slices were cut using a freezing stage microtome (Kryomat; Leitz, Germany). Ten series of sections were collected and stored in TBS/0.03% azide (100mM Tris, 15 mM NaCl, 0.5mM NaN_3_, pH 7.4) at 4°C.

Series of 40 μm-thick transverse sections of fixed spinal cord were immunolabelled as previously described. <u>Primary antibodies</u> (overnight) were: rabbit anti-PKCγ (1:500, sc-211, Santa Cruz Biotechnology); rabbit anti-serotonin (1:6000, #20080, Immunostar). Secondary antibodies (3 h) were: goat anti-rabbit IgG Alexa 488 (1:1000, Invitrogen) with DAPI (1:50,000, Sigma).

For BDA staining, free floating sections were incubated in 0.3% H_2_O_2_ and 10% methanol (30 mins). Sections were incubated in ABC vector (VectorLabs, UK) (30 mins) then amplified using biotinyl tyramide (1:75, PerkinElmer, USA), then left overnight at 4ºC with extra avidin FITC (1:500, #E2761, Sigma). Sections were washed between all steps. Sections were cover slipped with Mowiol.

CST axons were counted that crossed the midline, at two more lateral planes and at an oblique plane (Figure 3a) at C7 and C1. For each rat, we estimated the number of CST axons per cord segment by calculating the average number of CST axons per section and then multiplying by a scaling factor (number of sections cut per segment).

**Figure 3:**
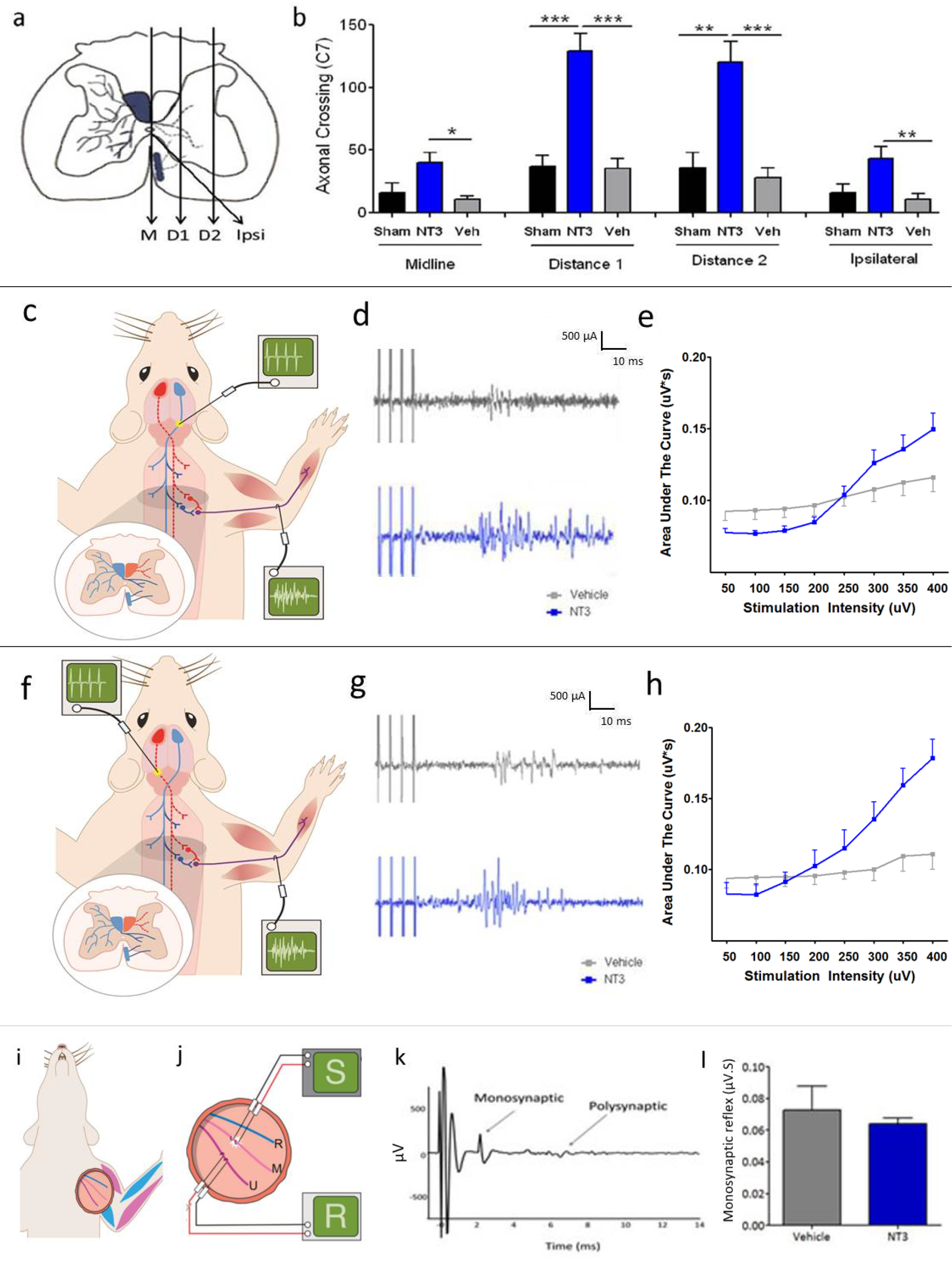
Delayed neurotrophin-3 treatment caused corticospinal tract sprouting and increased output in forelimb motor nerves during corticospinal tract stimulation. **a)** Corticospinal axons were anterogradely traced from the less-affected cortex and were counted at the midline (M), at two more lateral planes (D1 and D2) and crossing into grey matter from the ipsilateral, ventral tract (Ipsi). **b)** NT3 treatment caused an increase in the number of axons crossing at the midline (F_2,28_=6.1, p=0.007; *post hoc* p value=0.003), at two lateral planes denoted as D1 (F_2,28_=20.3, p<0.001, *post hoc* p value<0.001) and D2 (F_2,28_=13.8,p<0.001, *post hoc* p value<0.001) and from the ventral CST on the treated side (F_2,28_=5.2, p=0.013, *post hoc* p value=0.005). Although stroke by itself caused sprouting at the midline at C1 (planned comparison p=0.031), NT3 did not promote additional sprouting at C1. n=10/group were used for tract tracing. **c-e)** The CST from the less-affected hemisphere or **f-h)** lesioned hemisphere was stimulated in the medullary pyramids (before the decussation) and the motor output was recorded from the ulnar nerve on the treated side. **d, g)** The majority of spikes were detected between 17 ms and 45 ms, latencies consistent with polysynaptic transmission, in both vehicle-treated rats (grey) and NT3 treated rats (blue) when the less-affected or affected CST was stimulated. **e, h)** Stimulus intensity was increased incrementally from 50 μA to 400 μA and the area under the curve were measured (between 17 and 45 ms) after stimulation of the affected or less-affected hemisphere. NT3 treatment caused increased output in the ulnar nerve during stimulation of either the affected CST (two-way RM ANOVA intensity* group interaction F_1,14_=7.9, p=0.008, n=6 Vehicle, 6 NT3) or less affected CST (F_2,17_=9.8, p=0.01, n=4 Vehicle, n=6 NT3). **i, j)** The heteronymous reflex from median afferents to ulnar motor neurons was recorded in the axilla. **k)**The monosynaptic component was measured. **l)** NT3 did not increase the strength of the monosynaptic component.

The total length of serotonergic processes was measured using a standard method designed specifically to measure serotonergic sprouting after neurotrophin treatment (see refs in^17^) and which is well suited for quantification of dense terminal arbors (*e.g*., in the dorsal horn of the spinal cord). Processes were identified using the “adjust threshold” function in ImageJ and fiber lengths were measured in three areas: the dorsal horn, intermediate grey and ventral horn (Fig. 4a) in 3 sections per rat. We calculated the ratio of the sides ipsilateral and contralateral to NT3 treatment for the three areas separately.

**Figure 4:**
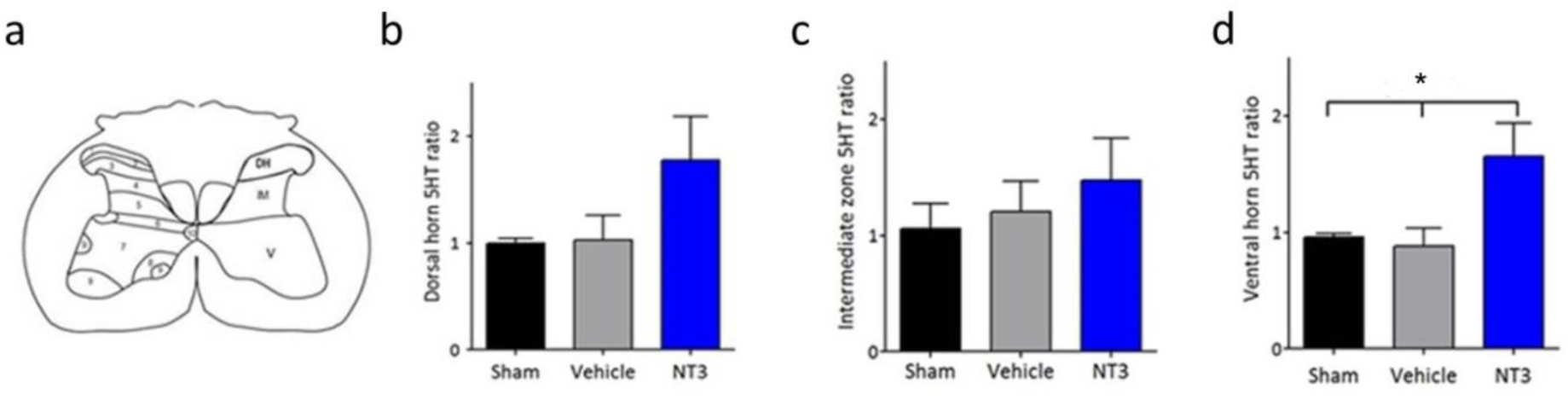
NT3 treatment caused plasticity of serotonergic axons in the C7 spinal cord. **a)** The total length of serotonergic arbors was measured in three regions shown in the schematic (DH, dorsal horn; IM, intermediate horn; V, ventral horn) on the treated side relative to the opposite hemicord. **b)** NT3 treatment did not increase serotonergic arbor length in the dorsal horn or **c)** the intermediate zone but **d)** did increase serotonergic arbor length in the ventral horn (one-way ANOVA F_2,22_=3.5, p=0.05; *post hoc* t-test p=0.025).

Immunofluorescence was visualized under a Zeiss Imager.Z1 microscope or a confocal Zeiss LSM 700 laser scanning microscope. Photographs were taken using the Axio Cam and AxioVision LE Rel. 4.2 or the LSM Image Browser software for image analysis.

### Radiolabelling of NT3

NT3 protein was radiolabelled with 50 μCi (1.85 MBq) N-succinimidyl [2,3-^3^H]propionate (^3^HNSP) and separated from unbound ^3^H-NSP using an ÄKTAprime purification system using a modification of a previous method^45,46^. 50 μg [^3^H]NT3 was injected intravenously as a 300 μL bolus into normal adult C57Bl/6 mice (7 to 8 weeks old, total n=25). Radiolabelled albumin was used as a baseline control because it does not cross the BBB efficiently. After 3, 15, 20, 30, 40, 60 or 90 min, brain, spinal cord and serum were taken for scintillation counting. The rate of influx (K_i_) was calculated from the Patlak plot^47^ of V_D_for [^3^H]NT3 and [^3^H]albumin against the plasma area under the curve.

The same treatment was repeated for mice 24 h after stroke, with an incubation period of 40 mins (n = 6). In this set of experiments, 0.074 MBq of ^14^C Sucrose (vascular marker) was injected towards the end of the incubation and the brain tissue samples also taken for capillary depletion analysis to distinguish NT3 or albumin in vascular endothelial cells from that in brain parenchyma. In brief, brain tissue was homogenized in physiological buffer (30 μl per mg of tissue) and 26% dextran (40 μl per mg of tissue) as described previously^48^. The homogenate was subjected to density gradient centrifugation (5,400 × g for 15 min at 4°C) to give an endothelial cell-enriched pellet and a supernatant containing the brain parenchyma and interstitial fluid (ISF). The homogenate, pellet and supernatant samples were solubilized and counted as described above. Distribution volume, V_D_, was calculated for all samples, including the endothelial pellet and brain parenchyma (ISF). The values were corrected for ^14^C Sucrose. Data was analysed for capillary fraction, parenchyma and whole brain using one way ANOVA and *post hoc* (Bonferroni) t-tests.

### Proteomic analysis

10 ug of protein was subjected to denaturing or non-denaturing SDS PAGE and visualised using Colloidal Coomassie brilliant blue staining. Each band was excised separately, digested enzymatically (with trypsin) and subjected to LC/MS/MS analysis (Dr. Steve Lynham, Proteomics Facility, KCL). In-gel reduction, alkylation and digestion with trypsin were performed prior to subsequent analysis by mass spectrometry. Cysteine residues were reduced with dithiothreitol and derivatised by treatment with iodoacetamide to form stable carbamidomethyl derivatives. Trypsin digestion was carried out overnight at room temperature after initial incubation at 37°C for 2 hours.

#### LC/MS/MS

Peptides were extracted from the gel pieces by a series of acetonitrile and aqueous washes. The extract was pooled with the initial supernatant and lyophilised. Each sample was then resuspended in 23L of 50mM ammonium bicarbonate and analysed by LC/MS/MS. Chromatographic separations were performed using an Ultimate LC system (Dionex, UK). Peptides were resolved by reversed phase chromatography on a 75 μm C18 PepMap column using a three step linear gradient of acetonitrile in 0.1% formic acid. The gradient was delivered to elute the peptides at a flow rate of 200 nL/min over 60 min. The eluate was ionised by electrospray ionisation using a Z-spray source fitted to a QTof-micro (Waters Corp.) operating under MassLynx v4.0. The instrument was run in automated data-dependent switching mode, selecting precursor ions based on their intensity for sequencing by collision-induced fragmentation. The MS/MS analyses were conducted using collision energy profiles that were chosen based on the mass-to-charge ratio (*m/z*) and the charge state of the peptide.

#### Database Searching

The mass spectral data was processed into peak lists using ProteinLynx Global Server v2.2.5 with the following parameters: (MS survey – No background subtraction, SG smoothing 2 iterations 3 channels, peaks centroided (top 80%) no de-isotoping; MS/MS – No background subtraction, SG smoothing 2 iterations 4 channels, peak centroiding (top 80%) no de-isotoping). The peak list was searched against the Uniprot database using Mascot software v2.2 using the following parameter specifications (Precursor ion mass tolerance 1.2 Da; Fragment ion mass tolerance 0.6 Da; Tryptic digest with up to three missed cleavages; Variable modifications: Acetyl (Protein N-term), Carbamidomethylation (C), Gln->pyro-glu (N-term Q) and Oxidation (M).

LC/MS/MS analysis and interrogation of the data against the Uniprot database identified NT3 from the excised and digested 1D gel bands. The results of the analysis and database searches are given in Supplementary Figure 8. Database generated files were uploaded into Scaffold 3 (v3.6) software (www.proteomesoftware.com) to create the .sfd file (PR320 LM 1D gel 12072012). All samples were aligned in this software for easier interpretation and used to validate MS/MS based peptide assignments and protein identifications. Peptide assignments were accepted if they contained at least two unique peptide assignments and were established at 100% identification probability by the Protein Prophet algorithm^49^. The result table includes probability scores (Mowse) for each peptide identified from the protein sequence. The Threshold Identity Score corresponds to a 5% chance of incorrect assignment. Peptides identified below these probabilities were accepted following manual inspection of the raw data to ensure that fragment ions correctly match the assigned sequence. The sequence coverage for each identified protein is represented in Supplementary Figure 8 in yellow highlights.

### Statistics

Statistical analyses were conducted using SPSS (version 18.0). Graphs show means ± SEMs (except where otherwise stated) and ‘n’ denotes number of rats. Asterisks (*,**,***) indicate p≤0.05, p≤0.01 and p≤0.001, respectively. Threshold for significance was 0.05. Histology and molecular biology data were assessed using Kruskal-Wallis and Mann-Whitney tests (due to small sample sizes). Serotonergic fibre lengths was analysed by region using one way ANOVA and *post hoc* (Bonferroni) t-tests. PKCγ data was analysed using Kruskal Wallis and Mann Whitney tests. Behavioural and MRI data were analysed using linear models and Restricted Maximum Likelihood estimation to accommodate data from rats with occasional missing values^50^. Akaike’s Information Criterion showed that the model with best fit for the horizontal ladder data had a compound symmetric covariance matrix, whereas for the sensory test and MRI data an unstructured covariance matrix was used. The model with best fit for the vertical cylinder had a compound symmetric covariance matrix, according to the –2 restricted log likelihood information criterion. Baseline scores were used as covariates. Degrees of freedom are reported to nearest integer. Normality was assessed using histograms. t-tests were two-tailed unless otherwise specified. Sample size calculations were presented previously^38^.

## Results

### MRI confirmed equivalent infarct volumes between stroke groups

Magnetic resonance imaging (MRI) confirmed that infarcts included the forelimb and hindlimb areas in sensorimotor cortex (Fig 1c). There was no difference in the mean infarct volume between stroke groups at 24 h, one or eight weeks after stroke (Fig. 1d). Loss of CST axons was assessed at 8 weeks in the upper cervical spinal cord using protein kinase C gamma (PKCγ) immunofluorescence^38,51^ (Fig. 1e). Stroke caused a 24% loss of CST axons in the dorsal columns relative to shams (Fig. 1f) with no difference between vehicle and NT3 treated rats. Together, the MRI and PKCγ histology data indicate that there were no confounding pre-treatment differences in mean infarct volumes and that NT3 did not act as a neuroprotective agent, as expected, based on our previous results^19^ and given that treatment was initiated after the majority of cell death will have occurred.

### Delayed NT3 improved forelimb sensory and motor function after stroke

We used the “adhesive patch” test to assess forepaw somatosensory function. A Sensory Score was obtained by attaching pairs of adhesive patches to each rat’s wrist on the dorsal side (Fig. 2a): a high score (*e.g*. 6) denotes that a rat preferentially removed the smaller stimulus from their less-affected wrist (*i.e*., did not first remove the larger stimulus on their affected wrist). The two stroke groups exhibited a similar lack of responsiveness to stimuli on their affected wrists after one week (Fig. 2b). Delayed treatment with NT3 caused recovery compared to vehicle: whereas vehicle-treated stroke rats showed a deficit relative to sham rats which persisted for eight weeks. Importantly, there were no confounding differences in the time taken to contact or to remove a patch from either their less-affected or affected paw: after stroke, NT3-treated rats and vehicle-treated rats took longer to contact an adhesive patch relative to shams, but there was no difference between NT3 and vehicle treated rats (Supplementary Fig. 1a). Moreover, neither stroke nor NT3 treatment caused any deficit in the additional time taken after contact to remove the patch (Supplementary Figure 1b). Thus, delayed treatment of disabled forelimb muscles with NT3 improved responsiveness to tactile stimuli after ischemic stroke.

Walking was assessed using a horizontal ladder with irregularly spaced rungs (Fig. 2c). Accurate paw placement during crossing requires proprioceptive feedback from muscle spindles^24^. After one week, the two stroke groups made a similar number of errors with their affected forelimb when crossing a horizontal ladder (Fig. 2d). Delayed NT3 treatment caused a progressive recovery after stroke whereas vehicle treated animals remained persistently impaired until the end of the study. This is consistent with previous work from our lab^17^,^19^. Stroke also caused a modest unilateral hindlimb impairment on the ladder; infusion of NT3 into the forelimb *triceps brachii* did not improve this (Supplementary Figure 2).

Neurotrophin-3 also restored the use of the affected forelimb for lateral support while rats reared in a vertical cylinder (Fig. 2e). After stroke and vehicle treatment, rats used their affected forelimb less often than shams. NT3-treated rats showed more frequent use of the affected forelimb relative to vehicle-treated rats (Fig. 2f).

We used force transducers to measure grip strength of each forelimb (Supplementary Fig. 3). Stroke caused transient weakness in both groups but infusion of NT3 into *triceps brachii* did not modify grip strength. We also found no evidence for pain (cold allodynia) on the affected or treated forelimbs, assessed by application of ice-cold acetone to the centre of the forepaw. Cold allodynia was induced neither by stroke nor NT3 treatment (Supplementary Fig. 4).

In summary, infusion of NT3 protein into the *triceps brachii* induced recovery on both sensory and motor tasks that require control of muscles by pathways including corticospinal pathways, serotonergic raphespinal pathways and proprioceptive circuits. Accordingly, we hypothesised that NT3 would induce neuroplasticity in multiple pathways.

### NT3 induced neuroplasticity in multiple spinal locomotor pathways

We examined anatomical neuroplasticity in the C7 cervical spinal cord because we knew from experiments using adult and elderly rats that the less-affected corticospinal tract sprouts at this level (as well as other levels) after injection of AAV-NT3 into muscles including *triceps brachii*^19^. Indeed, anterograde tracing from the less-affected hemisphere (Fig. 1b) revealed that infusion of NT3 protein increased sprouting of the CST in the C7 spinal cord (Fig. 3 a,b) across the midline and into the affected side at two more lateral planes, and also from the ventral CST.

We assessed neural output in the ulnar nerve on the affected side, whose motor neurons are also found in C7 (range: C4 to C8) that supply muscles in the forearm including the hand^42^,^52^. To do this, we recorded responses during electrical stimulation of either the spared less-affected corticospinal tract (Fig. 3c) or the partially-ablated corticospinal tract (Fig. 3f) in the medullary pyramids. NT3 treatment led to enhanced responses in the ulnar nerve during stimulation of the less-affected (Fig. 3 d, e) and more-affected (Fig. 3g, h) pathways. This result is consistent with the sprouting of traced CST axons (Fig. 3a,b) and indicates that CST axons from both the stroke hemisphere and the contralesional hemisphere formed new synapses and/or strengthened preexisting connections in the cord on the treated side, most likely on pre-motor interneurons that lie between CST axons and motoneurons^53,54^. However, we did not find any evidence that NT3 strengthened the short-latency reflex from afferents in the median nerve to motor neurons in the ulnar nerve (Fig. 3i-l). We also found that NT3 treatment caused serotonergic axons to sprout in the ventral C7 spinal cord (Fig. 4a-d). Anatomical and functional plasticity of corticospinal and raphespinal pathways is consistent with their expression of receptors for NT3^3,4,55–57^.

We conclude that NT3 caused neuroplasticity in multiple descending locomotor pathways including the raphespinal and the spared corticospinal tracts. These data are consistent with previous findings from our lab^17,19^ that peripherally-administered NT3 can, directly or indirectly, enhance supraspinal plasticity after stroke. Accordingly, next, we assessed the biodistribution of NT3 after peripheral administration.

### Neurotrophin-3 enters the PNS and CNS after peripheral administration

We measured the amount of total (rat and human) NT3 in the *triceps brachii* and C7 spinal cord. ELISA was performed using a subset of five rats per group withdrawn at random from the study at the four-week time point: this revealed an increase in total NT3 protein levels in the *triceps brachii* on the treated side (Fig. 5a) and, surprisingly, on the untreated side (perhaps due to NT3 in endothelial cells; see below). We were not able to detect any increase in total NT3 in the C7 spinal cords (Supplementary Fig. 5). However, ELISA cannot distinguish exogenous human NT3 from endogenous rat NT3 because the amino acid sequences for mature human and rat NT3 are identical^58,59^. Because ELISA did not allow us to detect any small increases in exogenous human NT3 against the background of endogenous rat NT3 in the CNS, we next used a more sensitive method for measuring trafficking of NT3 across the blood-CNS barrier.

**Figure 5:**
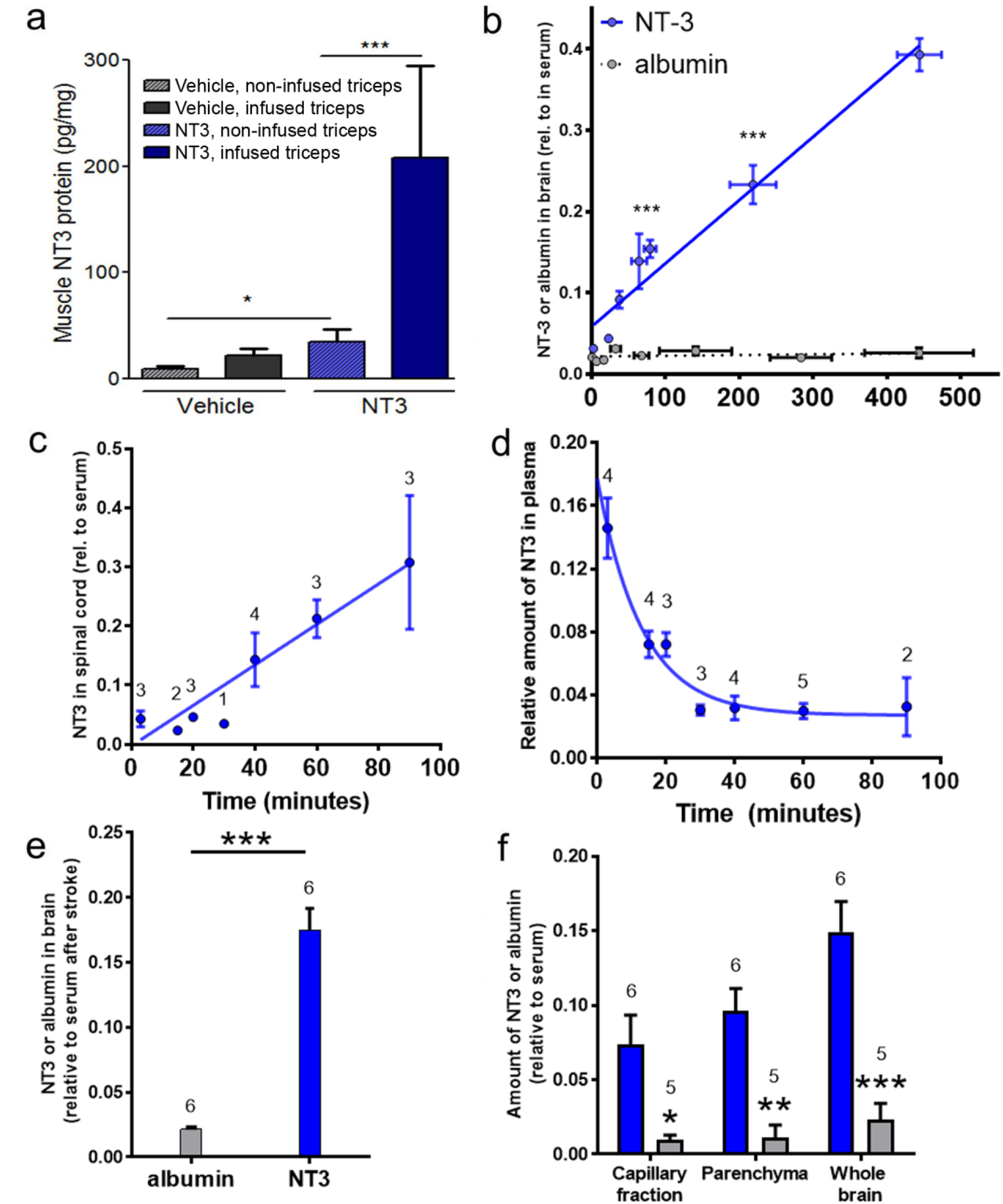
Levels of NT3 were increased in *triceps brachii*, in brain and in spinal cord. **a)** ELISA revealed that, in NT3-treated rats, there was an increase in NT3 in the infused *triceps brachii* (Mann-Whitney vs. vehicle, p=0.032) as well as in the non-infused *triceps brachii* (Mann-Whitney vs. vehicle, p=0.032). n=5 rats/group. **b)** [^3^H]NT3 entered the brain more abundantly than [^3^H]albumin. Regression of [^3^H]NT3 (blue circles) and [^3^H]albumin (grey circles) against time after *iv* injection (n=2–5 mice/time) in adult mice. T-tests for NT3 vs albumin all significant for incubation times of 3, 15, 20 30, 40 and 60 (p values from <0.01 to <0.0001 ***). The Volume of Distribution of [^3^H] NT3 or [^3^H] albumin in brain (V_D_=A_m_/C_P_) is calculated as a ratio of counts per minute (CPM) in 1 μg of brain and CPM in 1 μl of serum for each time point and plotted against exposure time given by the term ∫_0_^t^Cp(τ)dτ/Cp. The rate of influx (K_i_) was calculated from the Patlak plot^47^ of V_D_for [^3^H]NT3 and [^3^H]albumin against the plasma area under the curve. At 1.3 × 10^-5^ μl/mg/s, the unidirectional influx constant for NT3, K_i_, is 100 times that of the albumin 1.3 ×10^-7^ μl/mg/s). **c)**[^3^H]NT3 entered the cervical spinal cord more abundantly than [^3^H]albumin. **d)**Plasma half-life of NT3 for the normal adult mouse is ∼10 min (estimated from 1/normalised serum values). **e)** Twenty-four hours after cortical ischemia, [^3^H]NT3 entered the brain more abundantly than [^3^H]albumin, measured 40 minutes after *iv* injection. **f)**Twenty-four hours after cortical ischemia, levels of[^3^H]NT3 were higher in parenchyma depleted of endothelial cells than levels of [^3^H]albumin 40 minutes after *iv* injection (parenchyma p < 0.01; capillary fraction p < 0.05; and whole brain P < 0.001, two-way ANOVA with Bonferroni *post hoc* tests). Results were corrected for the value of V_D_ for ^14^C Sucrose. Number of mice per group stated on panels.

Recombinant NT3 protein was radiolabelled and purified. [^3^H]NT3 was injected intravenously into adult mice. Radiolabelled albumin was used as a control because it does not enter the CNS efficiently from the bloodstream. After 3, 15, 20, 30, 40, 60 or 90 minutes, brain, spinal cord and serum were taken for scintillation counting. NT3 progressively accumulated in the intact brain (Fig. 5b) and cervical spinal cord (Fig. 5c). In plasma, the half-life of NT3 was short (Fig. 5d). Our data is consistent with that from others who have shown that radiolabelled NT3 rapidly crosses the barriers between the blood and an intact CNS^26–28^ and that a small amount of intact NT3 accumulates in the brain and cervical spinal cord (although the majority of NT3 is cleared rapidly from the bloodstream)^26–28^. For example, after injection of NT3 into the brachial vein (which provides drainage from the *triceps brachii*), NT3 accumulates in the cortex, striatum, brainstem, cerebellum, sciatic nerve (and other regions of the nervous system involved with locomotion)^27^.

Next, [^3^H]NT3 or [^3^H]albumin was injected intravenously into adult mice 24 hours after cortical ischemia. 40 minutes later, tissues were taken for scintillation counting. In contrast to [^3^H]albumin, [^3^H]NT3 accumulated in the brain (Fig 5e). To confirm entry of [^3^H]NT3 into brain parenchyma beyond endothelial cells, capillaries were depleted by gradient centrifugation to yield a supernatant containing brain parenchyma and an endothelial cell-enriched pellet^60^. [^3^H]NT3 entered parenchyma (depleted of endothelial cells) at a level above that seen for [^3^H]albumin (Fig. 5f). Transport of NT3 into the CNS is apparently a receptor-mediated process^61^ as shown by 1) the expression of NT3 receptors in rodent and human CNS capillaries^62,63^ and 2) the ability of non-radiolabelled NT3 to compete for uptake of radiolabelled NT3 into the CNS^12,13,44^.

In addition, we and others have shown that NT3 enters the PNS after peripheral administration: after intramuscular overexpression of AAV encoding NT3, NT3 levels are elevated in the blood stream and NT3 accumulates in the ipsilateral DRG^17,19^. We also found some evidence that NT3 is retrogradely transported from muscle to ipsilateral motor neurons^17^. This is consistent with data showing that 1) the blood-nerve barrier in DRGs is permeable to proteins like NT3^64^, 2) that after intravenous injection, radiolabelled NT3 accumulates in the sciatic nerve^27^ and 3) that NT3 is retrogradely transported from muscle to the spinal cord or DRG in nerves^5,7,65,66^. We conclude that neuroplasticity occurred in multiple locomotor pathways because peripherally-administered NT3 bound to receptors in the PNS and CNS.

### BOLD signal in perilesional somatosensory cortex evoked by stimulation of the affected wrist recovers with time after stroke and is not further enhanced by NT3

To explore the mechanism whereby NT3 improved responsiveness to stimuli attached to the affected wrist (Fig. 2b), we performed functional brain imaging (BOLD-fMRI) during low threshold (non-noxious) intensity electrical stimulation of the affected wrist (Supplementary Fig. 6). As expected, prior to stroke, stimulation of the wrist resulted in a higher probability of activation of the opposite somatosensory cortex (Supplementary Fig. 6a). fMRI performed one week after stroke confirmed that somatosensory cortex was not active when the affected paw was stimulated in either vehicle or NT3 treated rats (p>0.05, Supplementary Fig. 6b). This supports our claim above that there were no early differences between groups that could be explained by neuroprotection. fMRI performed eight weeks after stroke revealed a trend towards perilesional re-activation of somatosensory cortex in both vehicle and NT3 treated groups (p<0.05, Supplementary Fig. 6c). This is in line with human brain imaging studies showing that spontaneous sensory recovery is increased after stroke when more-normal activity patterns are observed on the affected side of the brain^67^. However, these probabilities of re-activation were not big enough to survive correction for testing of multiple voxels (p-values>0.01) although clearly they are in a location that might mediate recovery of somatosensation.

A longitudinal analysis showed that at 8 weeks (relative to pre-stroke baseline), there was some evidence that rats treated with Neurotrophin-3 showed increased probability of activation of perilesional cortex (Supplementary Fig. 6d, p<0.05) and showed decreased probability of activation of somatosensory cortex on the less-affected hemisphere (p<0.05) relative to vehicle-treated stroke rats. However, these apparent differences did not survive correction for testing of multiple voxels (p<0.01). We conclude that both groups showed partial, spontaneous restoration of more-normal patterns of somatosensory cortex activation^68,69^ but, conservatively, that NT3 did not further increase probability of activation of any supraspinal areas. These conclusions are consistent with previous fMRI data from our laboratory^19^; we propose that the additional recovery of somatosensory function after NT3 treatment (Fig. 2b) is due to changes in the spinal cord rather than in supraspinal areas.

### Proteomic analysis of NT3

The batch of neurotrophin-3 protein that we used was produced more than a decade ago by Genentech. We sought to determine whether any degradation had occurred and to confirm its amino acid sequence so that identical preparations of neurotrophin-3 could be made for future experiments. Supplementary Figure 7 depicts results from a non-denaturing gel showing a higher molecular weight band (30 kDa) and a lower molecular weight band (14 kDa) consistent, respectively, with dimeric mature NT3 and monomeric mature NT3. There was no evidence of degradation or aggregation. Each band was excised separately, digested enzymatically (with trypsin) and subjected to LC/MS/MS analysis. Proteomic analysis was consistent with both bands being mature NT3 with no evidence of residual prepro sequences (Supplementary Figure 8). We conclude that the higher molecular weight band is not preproNT3 (∼30 kDa) but rather corresponds to dimeric mature NT3. This facilitated our ongoing experiments to evaluate NT3 as a therapy for stroke because most commercial preparations of NT3 consist of mature NT3 rather than preproNT3.

## Discussion

Treatment of disabled arm muscles with NT3 protein, initiated 24 hours after stroke, caused changes in multiple locomotor circuits, and promoted a progressive recovery of sensory and motor function in rats. The fact that NT3 can reverse disability when treatment is initiated 24 hours after stroke is exciting because the vast majority of stroke victims are diagnosed within this time frame^2^. In contrast, the gold-standard drug for ischemic stroke, tPA, needs to be given within a few hours and is only administered to a minority. Thus, NT3 could potentially be used to treat an enormous number of victims.

NT3 has good clinical potential. Firstly, Phase II clinical trials show that doses up to 150 μg/kg/day are well tolerated and safe in healthy humans and in humans with other conditions^29,31–33^. We used a threefold lower dose (48 μg/kg/day) in this study: in future experiments we will optimize the dose and duration of treatment because it is possible that a higher dose of NT3 would promote additional recovery after stroke. Secondly, there is good conservation from rodents to primates including humans in the expression of receptors for NT3 in the locomotor system^3,4,70–72^. Thirdly, in none of our rodent experiments has NT3 treatment caused any detectable pain, spasticity or muscle weakness (in line with the human trials); rather, after bilateral corticospinal tract injury in rats, intramuscular delivery of AAV-NT3 reduced spasticity, slightly improved grip strength and showed a trend towards reducing mechanical hyperalgesia^17^.

In this study and in a previous study we used functional MRI combined with electrical stimulation of the wrist in an effort to discover what neuroplasticity underlies recovery of somatosensory responsiveness to adhesive patches attached to the wrist. We confirmed work by others that recovery correlated well with more-normal patterns of increased BOLD signal surrounding the infarct (potentially in spared somatosensory cortex)^68,69,73,74^ but we did not find strong evidence that NT3 further increased peri-lesional (or other) activation (either in this study or in our previous study^19^). Instead, we now propose that NT3 increased somatosensory recovery by inducing neuroplasticity in spinal circuits involving cutaneous afferents. This is plausible because cutaneous afferents which mediate tactile sensitivity express TrkC receptors^6^. Moreover, others have shown that dl3 spinal interneurons can gate cutaneous transmission^75^. We have previously shown that NT3 normalises post-activation depression of output from spinal circuits evoked by stimulation of low threshold afferents from the treated wrist^17^ (which might include cutaneous afferents as well as proprioceptive afferents) although in those experiments we measured motor output rather than sensory transmission. In the future one might examine whether NT3 modulates gating of somatosensory inputs from the wrist to spinal interneurons^75^. However, in the present work, the deficits in somatosensory responses were modest and might be difficult to dissect.

With regard to corticospinal neuroplasticity, we have shown twice previously (in adult and elderly rats) that the less-affected corticospinal tract sprouts across the cervical midline after injection of AAV-NT3 into affected forelimb muscles^19^. Others have shown that intrathecal infusion of NT3 induces sprouting of the corticospinal tracts^16^ and that injection of vectors encoding NT3 into muscles^10^ or nerve^11^ can induce corticospinal tract sprouting. Here, our anatomical tracing confirmed that the less-affected corticospinal tract sprouted after infusion of NT3 protein into triceps and in future we will trace both tracts. This is because, in the present study, neurophysiology revealed that both corticospinal tracts underwent plasticity after unilateral infusion of NT3 protein. We propose that spared CST axons sprouted after NT3 entered the CNS from the systemic circulation. This is consistent with data from us and others showing that radiolabelled NT3 entered the brain and spinal cord after intravenous injection^26–28^. Moreover, it has been shown that endogenous muscle spindle-derived cues induce sprouting of descending pathways after spinal cord injury in adult mice^24^; given that muscle spindles make NT3 endogenously^25^, it is plausible that infusion of supplementary NT3 to muscle might enhance corticospinal sprouting after stroke.

It is also notable that infusion of NT3 into a proximal forelimb extensor improved the accuracy of use of the affected forelimb when walking on a horizontal ladder but did not improve the accuracy of use of the affected hindlimb; this implies that circulating NT3 is not sufficient to improve hindlimb movements. Moreover, we did not find any evidence that NT3 strengthened the short-latency reflex between afferents in the median nerve and motor neurons in the ulnar nerve; this may be because we infused NT3 into the *triceps brachii* whose afferents do not run in the median or ulnar nerve. This is consistent with previous work of ours showing that a reflex may be strengthened when its afferent comes from a muscle expressing higher levels of NT3 but not when its afferent comes from a muscle lacking transgenic expression of NT3^17^. Finally, infusion of NT3 protein into *triceps brachii* did not improve forelimb grip strength: however, the grip strength task probably depends more on strength in hand and digit flexor muscles (into which NT3 was not infused) than on *triceps brachii* (elbow extensors). Indeed, in previous work, injection of AAV-NT3 into proximal and distal flexor muscles did modestly improve grip strength^17^. Taken together, these results indicate that it may be important to target NT3 to multiple muscles.

However, it is not straightforward to reconcile all our findings with a single mechanistic explanation. It is possible that, additionally, NT3 was trafficked from *triceps brachii* in axons to motor neurons and/or by DRG neurons where it induced expression of a molecule that was secreted and induced CST sprouting (e.g., BDNF or IGF1^76,77^). NT3 is certainly trafficked to ipsilateral motor neurons and DRG after intramuscular delivery^5,7,17,19,65^ and in this study we also showed, unexpectedly, a small increase in contralateral *triceps* (perhaps from NT3 in endothelial cells). Diffusion of NT3 within neuropil is inefficient^78^ but spinal motor neuron dendritic arbors can be very large; some even extend across the midline^79^ and these might provide a widespread source of cues for supraspinal axonal plasticity (*e.g*., across the midline). To seek DRG-secreted factors, we have performed RNAseq of cervical DRG after injection of AAV-NT3 into forelimb flexors^17^,^80^. In the future we will also seek motor neuron-derived cues.

Finally, it is interesting that the recovery continues even after infusion of NT3 is discontinued at four weeks. This is encouraging, from a translational perspective. We propose that the four-week long NT3 treatment induces changes in target neurons that persist (*e.g*., due to sustained modifications in gene expression). Indeed, longer treatment with NT3 induces different intracellular signalling events in sensory axons than does brief treatment, thereby enhancing terminal branching^81^. In the future, we will seek factors that are persistently increased in target neurons after NT3 treatment is discontinued. Additionally, it may be that NT3 induces sprouting of CST axons that (after cessation of treatment) is followed by selection of synapses (e.g., strengthening or pruning) by a mechanism that is independent of NT3. For example, it is known that corticomotoneuronal axon synapses are pruned by repulsive PlexinA1-Sema6D interactions^82^. To begin to dissect the mechanisms whereby NT3 promotes neuroplasticity and recovery after peripheral delivery, we are setting up a mouse model of stroke.

In summary, treatment of disabled arm muscles with NT3 (initiated in a clinically-feasible time-frame) induces multilevel spinal and supraspinal neuroplasticity, improves walking and reverses a tactile sensory impairment.

## Acknowledgments

Genentech provided neurotrophin-3 protein under Materials Transfer Agreement. Thanks to Dr Joe Lewcock for advice. Thanks to Dr Owen O’Daly for help with SPM. Thanks to members of the Moon Lab for comments on the manuscript. The research leading to these results has received funding from the European Research Council under the European Union’s Seventh Framework Programme (FP/2007–2013) / ERC Grant Agreement n. 309731 as well as by a Research Councils UK Academic Fellowship and by the Medical Research Council (MRC), the International Spinal Research Trust and the British Pharmacological Society (BPS)’s Integrative Pharmacology Fund. This study was also supported by the Dowager Countess Eleanor Peel Trust and a Capacity Building Award in Integrative Mammalian Biology funded by the Biotechnology and Biological Sciences Research Council, BPS, Higher Education Funding Council for England, Knowledge Transfer Partnerships, MRC and Scottish Funding Council.

## Conflicts of Interest

There are no conflicts of interests of the authors.

## Author contributions

D.D and L.M designed the experiments and wrote the manuscript. D.D and L.M. performed the rat experiments. K.B. and S.M. assisted with neurophysiology. S.W., D.C., C.S., M.B. and T.W. assisted with MRI scans and analysis. S.D., D.D. and D.B. performed mouse NT3 uptake experiments. Q.C. and H.D.S. performed ELISA. All authors provided material and commentary for the manuscript.

Correspondence and requests for materials should be addressed to L.M

## Supplementary Information

**Supplementary Figure 1:**
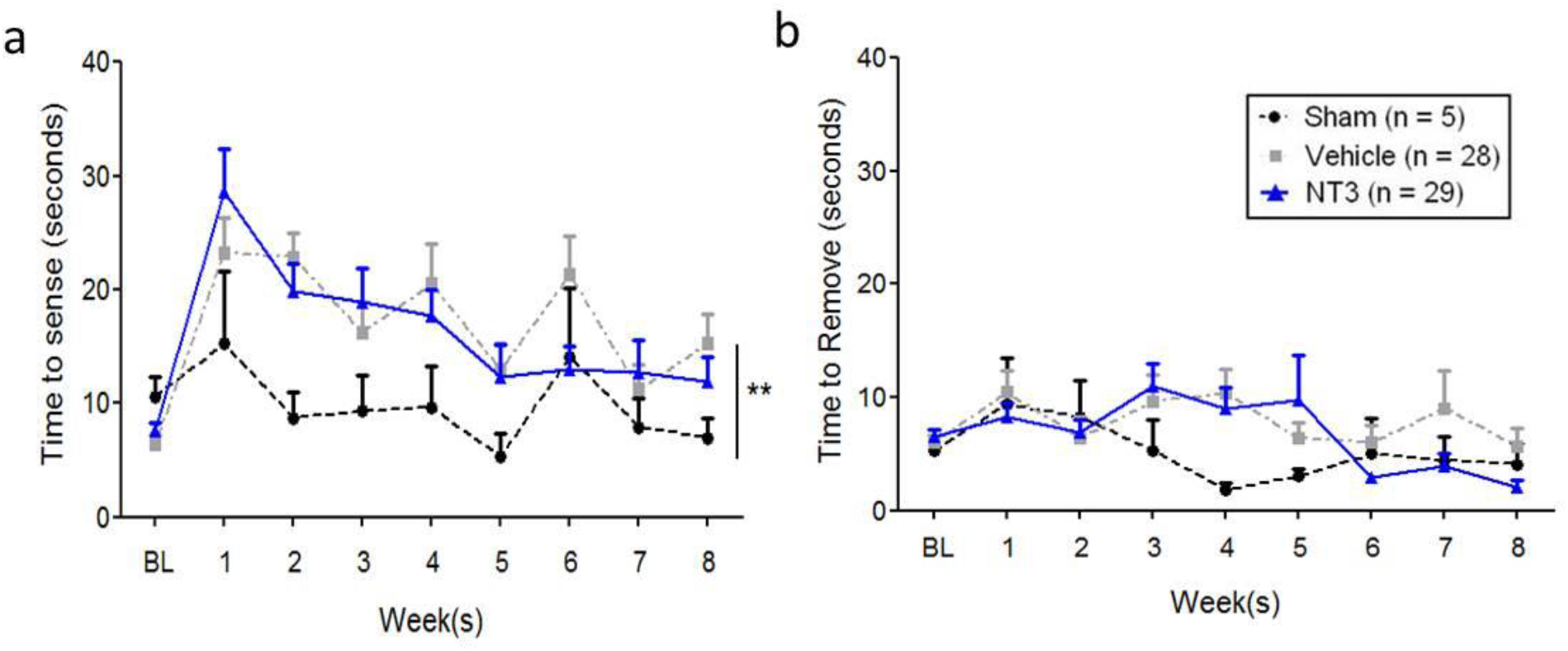
Stroke caused a transient impairment in responding to tactile stimuli placed on the affected wrist. **a)** After stroke, NT3-treated rats and vehicle-treated rats took longer to contact the adhesive patch relative to shams (linear model; F_2,54_=5.2, p=0.008, *post hoc* p values=0.005 and 0.002), and there was a similar amount of spontaneous recovery with time in both NT3 and vehicle treated groups (NT3 *vs*. vehicle, *p*=0.55). **b)** Stroke did not cause any deficit in the additional time taken after contact to remove the first patch (linear model; F2,55 = 0.95, p = 0.39) and there was no additional effect of NT3 (NT3 *vs*. vehicle, *p*=0.36).

**Supplementary Figure 2:**
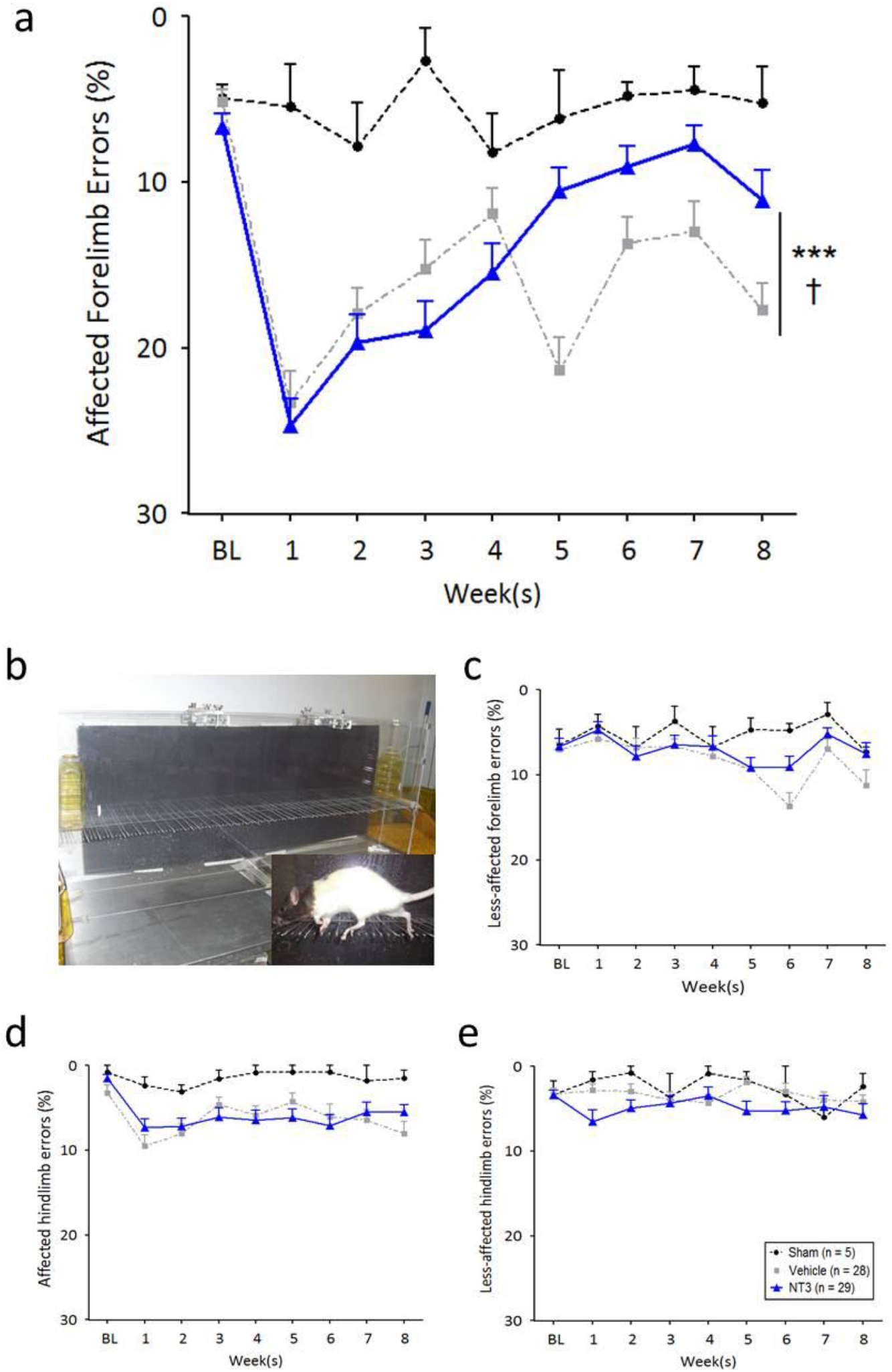
Focal cortical stroke caused impairment of the affected forelimb but modest or no impairment of the three other limbs. **a)** After stroke, NT3 treated rats recovered function of their affected forelimb on the ladder test relative to stroke vehicle controls and sham rats. NT3 treated rats recovered fully relative to shams (linear model and t-tests, p≤0.05). *** denotes group difference, p <0.05; † denotes interaction of group with time, p<.05. This subpanel is reproduced from Figure 2 to allow comparison with other subpanels). **b)** shows photograph of the horizontal ladder set up and insert shows a rat traversing the ladder. **c)** There was no difference in the number of foot faults made in any of the groups using the less affected forelimb, **(d)** the contralateral hindlimb or **(e)** the ipsilateral hindlimb, p-values>0.05. Linear models.

**Supplementary Figure 3.**
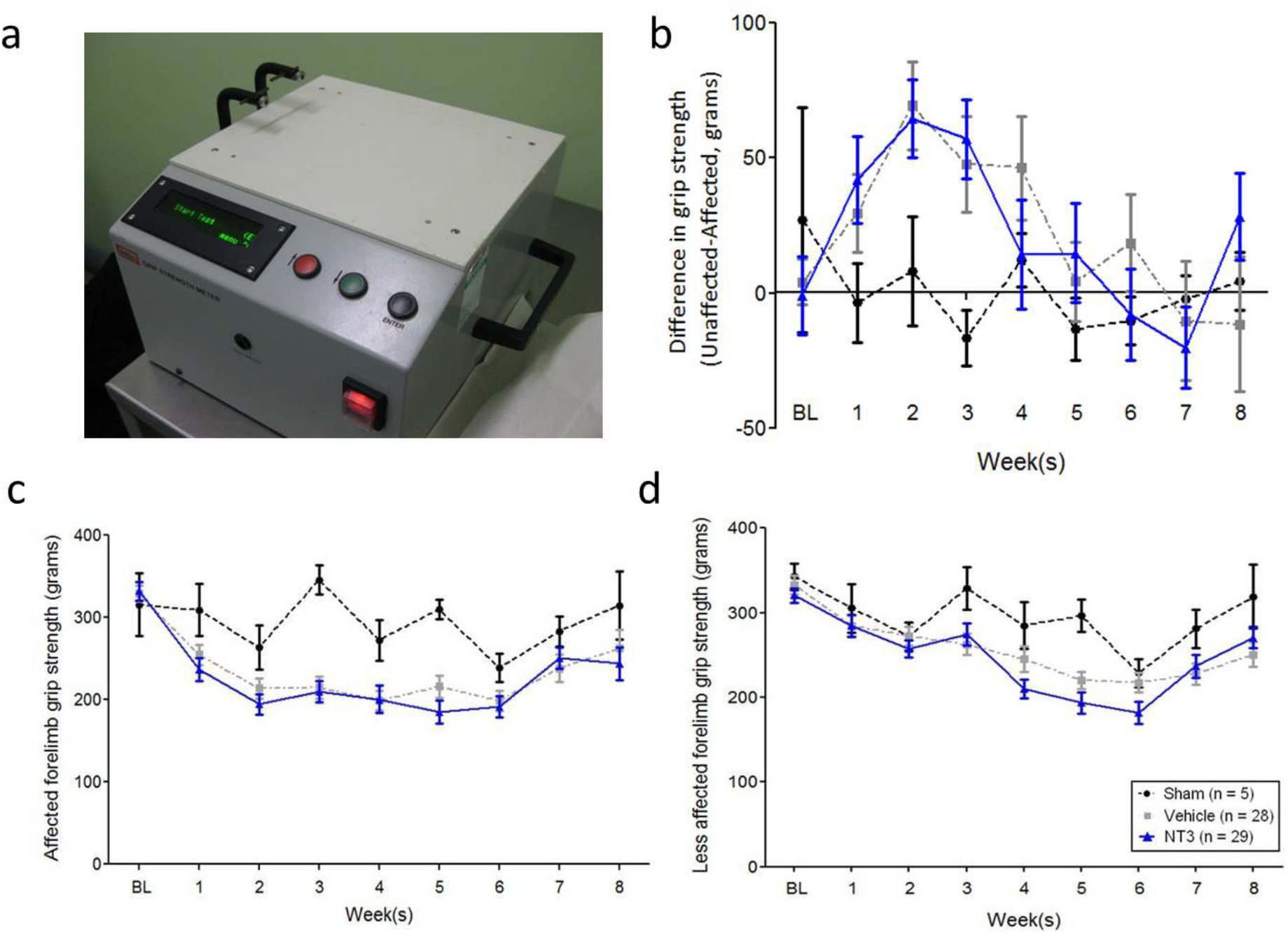
Focal cortical stroke only transiently affected upper limb strength, and NT3 neither strengthened nor weakened upper arm strength. **a)** The testing apparatus. The rats held bilaterally on to the pair of force transducers (top left) and the rat was pulled away horizontally and perpendicularly (towards the right) until the bilateral grip was broken. The force transducers provide a measure of strength (grams) for each upper limb. An average of three trials was taken per rat per week. Grip strength (grams) are presented as group means ± SEMs. **b)**Grip strength of the affected limb was subtracted from the unaffected limb strength, as an internal control (e.g., to control for differences in motivation, etc.). **c)** Grip strength for the affected limb. **d)** Grip strength for the less-affected limb. Stroke caused a weak trend towards a transient decrease in strength on the limb affected by stroke (relative to shams; time F7,349=1.8, p=0.09; p-values p=0.121 and 0.124, respectively) but multiple pairwise comparisons did not show significant differences at any timepoint (all p>0.05). There was no difference between NT3 and vehicle treated rats overall(group F_2,51_=1.3, p=0.27; group × time F14,351=0.72), or at any time point (all p values>0.05).

**Supplementary Figure 4:**
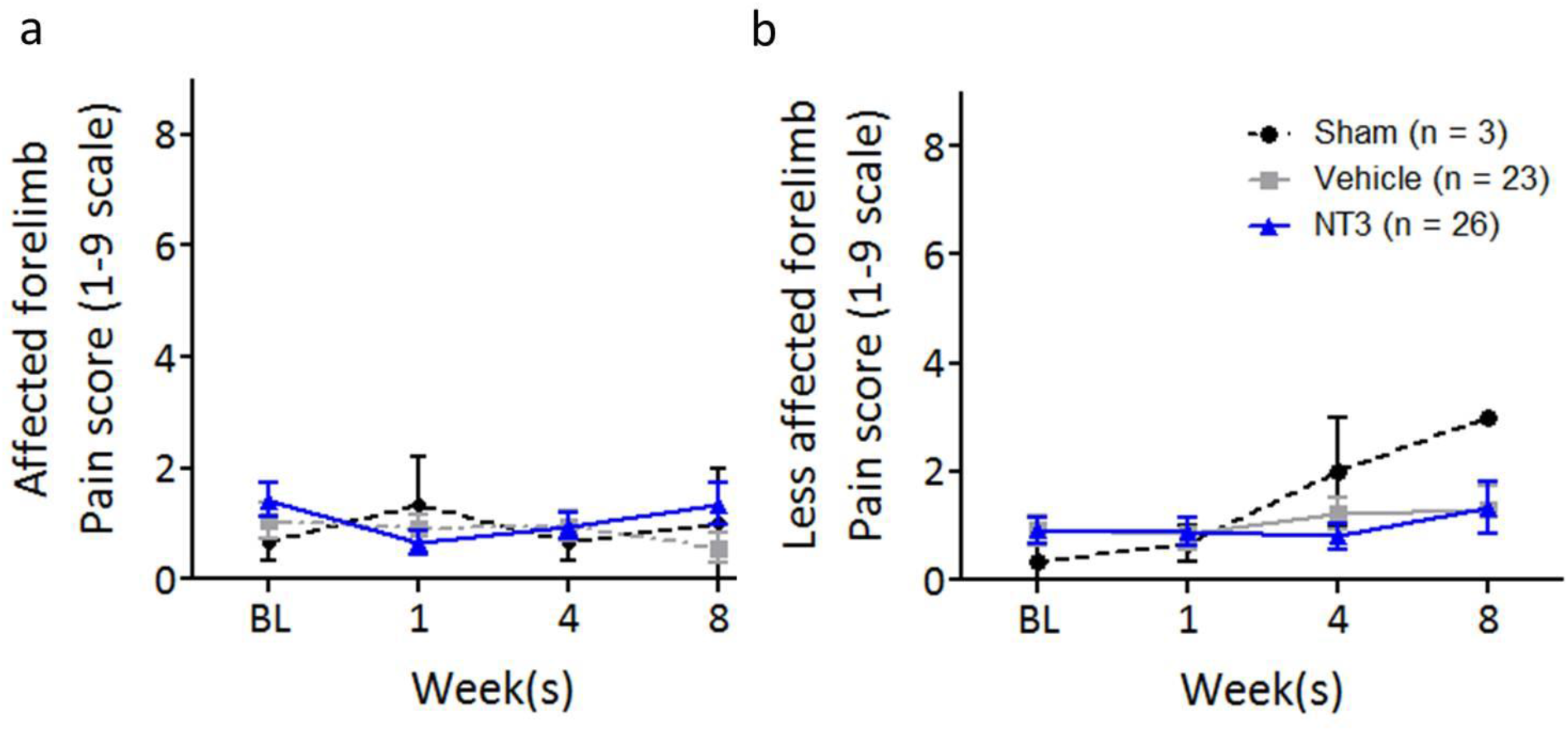
Cold allodynia was caused neither by focal cortical stroke nor by treatment with neurotrophin-3. The acetone test was used to see whether stroke and/or NT3 treatment caused any change in cold allodynia pain responses. The test involves applying a drop of acetone to the **a)** affected or **b)** less affected forelimb, and then allocating a score between 0 and 9: higher numbers denote a heightened pain response. There is no evidence of painful behaviour based on this test in either forelimb. RM ANCOVA with Bonferroni post hoc tests.

**Supplementary Figure 5:**
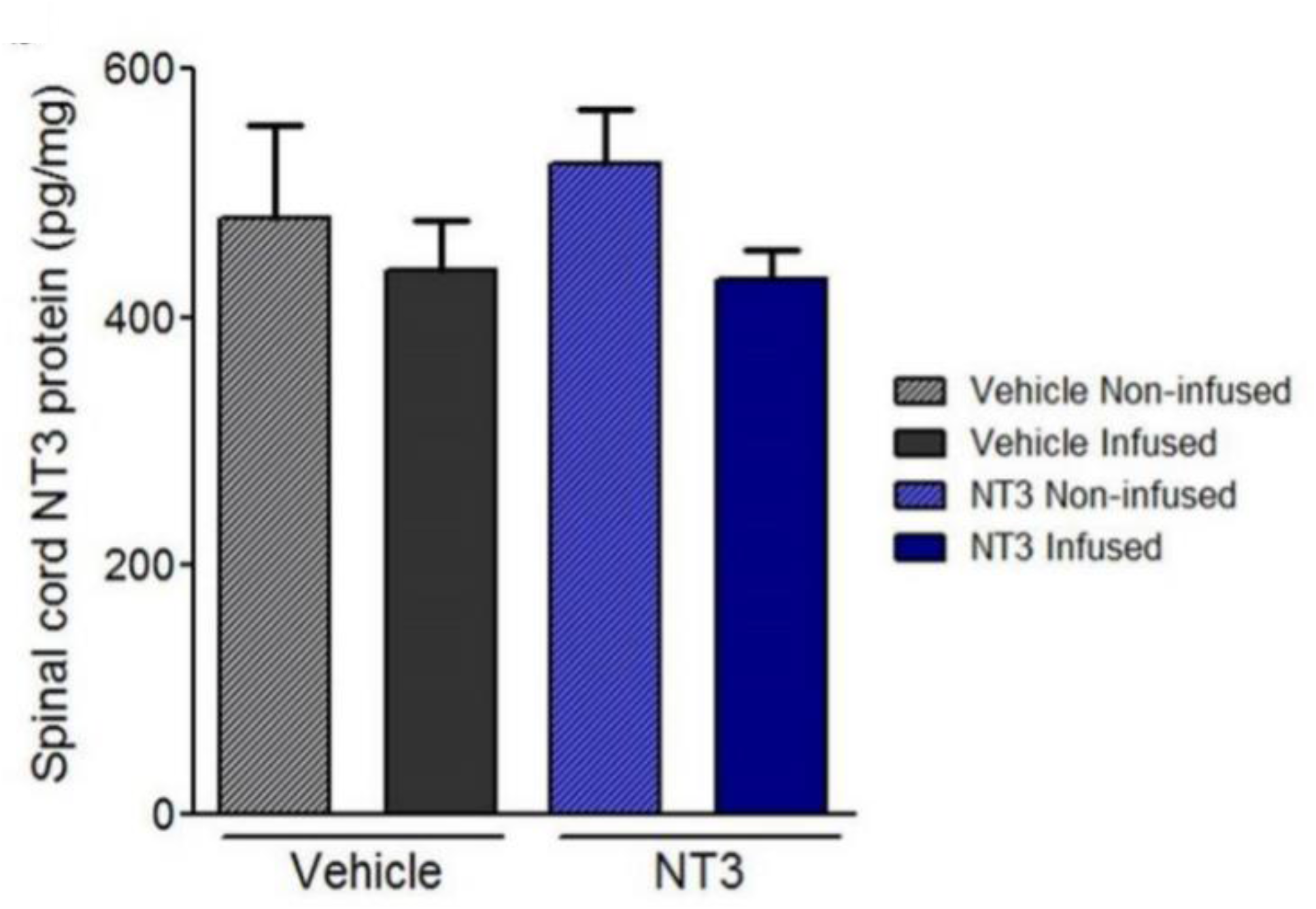
ELISA revealed that infusion of NT3 into *triceps brachii* did not cause detectable elevation of NT3 in homogenates of cervical spinal cord hemicords on the infused or non-infused side of the body (Mann Whitney p-values=0.84, 0.42, respectively).

**Supplementary Figure 6:**
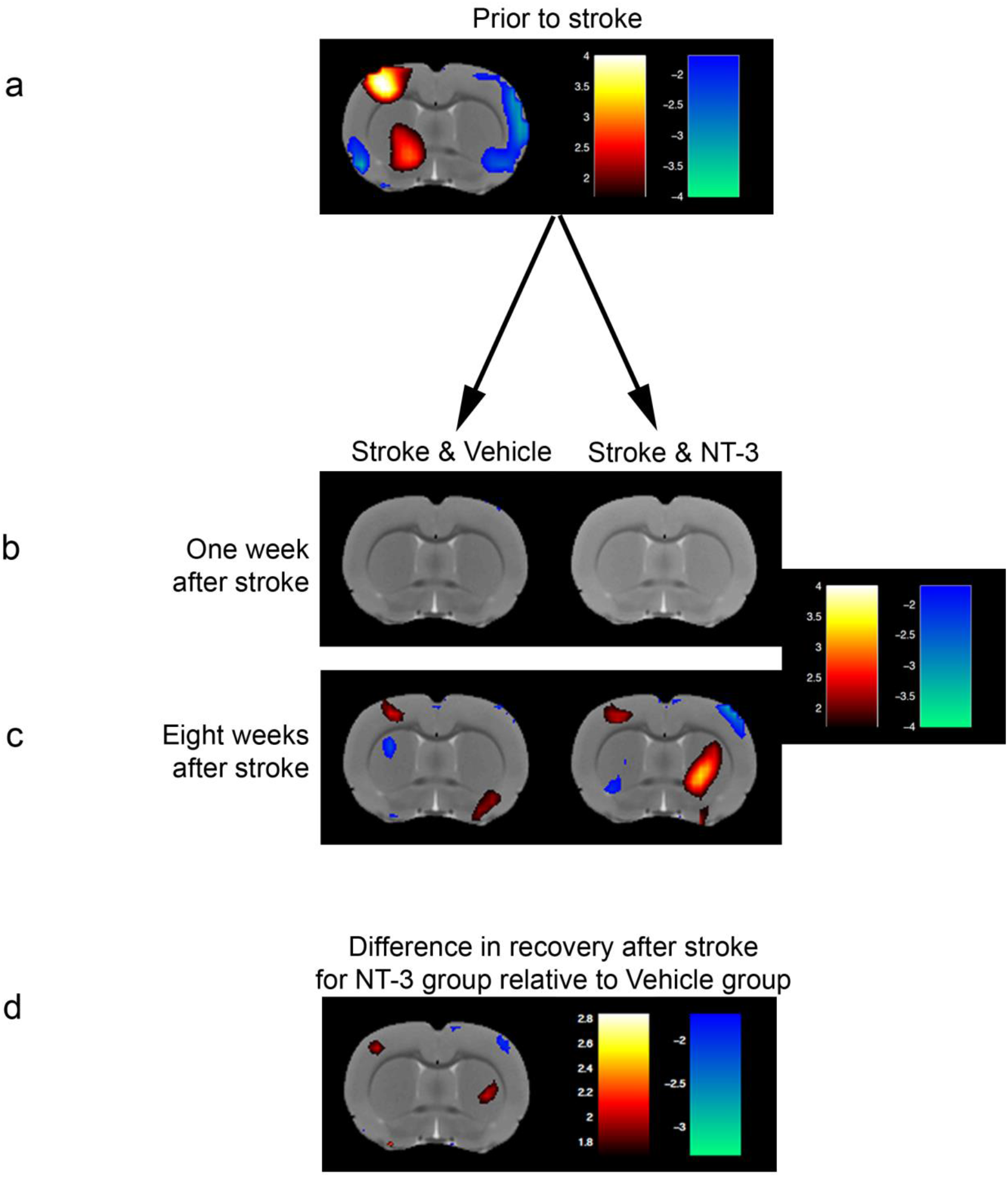
Functional brain imaging during stimulation of the affected wrist revealed no enhanced probability of perilesional activation by neurotrophin-3. The same rats were imaged prior to stroke and then one week and eight weeks after stroke and intramuscular treatment with either NT3 or vehicle. Scans with obvious imaging artefacts were discarded, leaving final group numbers of n=7, 9, 7 and n=9, 8, 4 at weeks 0, 1 and 8 for NT3 and vehicle treated groups respectively. Red voxels denote greater probability of activation during stimulation (versus stimulation off) whereas blue voxels denote lesser probability of activation during stimulation (versus stimulation off). **a)** Prior to stroke, stimulation of the dominant paw led to a strong probability of activation in the opposite somatosensory cortex. **b)** One week after stroke, this activation was abolished by infarction. **c)** Eight weeks after stroke, there was a slight trend towards a small perilesional area of reactivation in both groups. **d)** There was a slight trend towards greater perilesional reactivation in the NT3 group versus the vehicle group at 8 weeks (relative to their baselines). However, all these heat maps of groups of rats show t-values obtained by Statistical Parametric Map analysis without correction for multiple testing (p<0.05) and there were no differences between the two groups for any voxels when the threshold for significance was corrected for multiple testing (p<0.01; this data is not shown as the heat map was black). Red voxels denote greater probability of activation during stimulation for the NT3 group than for the vehicle group whereas blue voxels denote lesser probability of activation for the NT3 group than for the vehicle group. When stimulating the less-affected wrist, there were no differences between the two groups for any voxels when the threshold for significance was corrected for multiple testing (p<0.01; this data is not shown as the heat map was black).

**Supplementary Figure 7:**
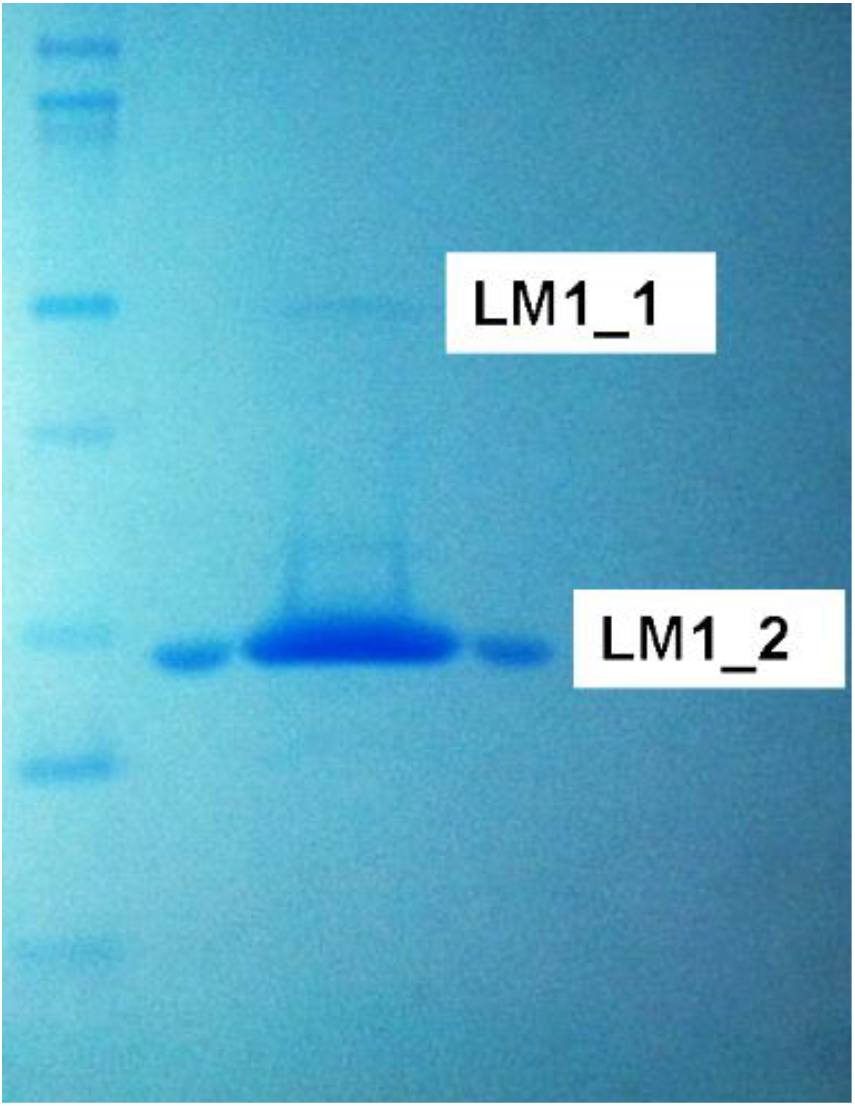
Different amounts of recombinant NT3 were run in four lanes of an SDS PAGE gel. Two sizes of band of interest (LM1_1 and LM1_2) were detected following staining with Colloidal Coomassie brilliant blue. These protein bands were excised prior to separate enzymatic digestion and LC/MS/MS analysis. The apparent molecular weight of the upper band (∼30kDa) is consistent with either the pro-Neurotrophin-3 precursor form or a dimer of the mature NT3 protein, whilst the lower band (∼14kDa) is consistent with the mature NT3 protein. Sequencing revealed that both bands represent the mature NT3 protein.

**Supplementary Figure 8:**
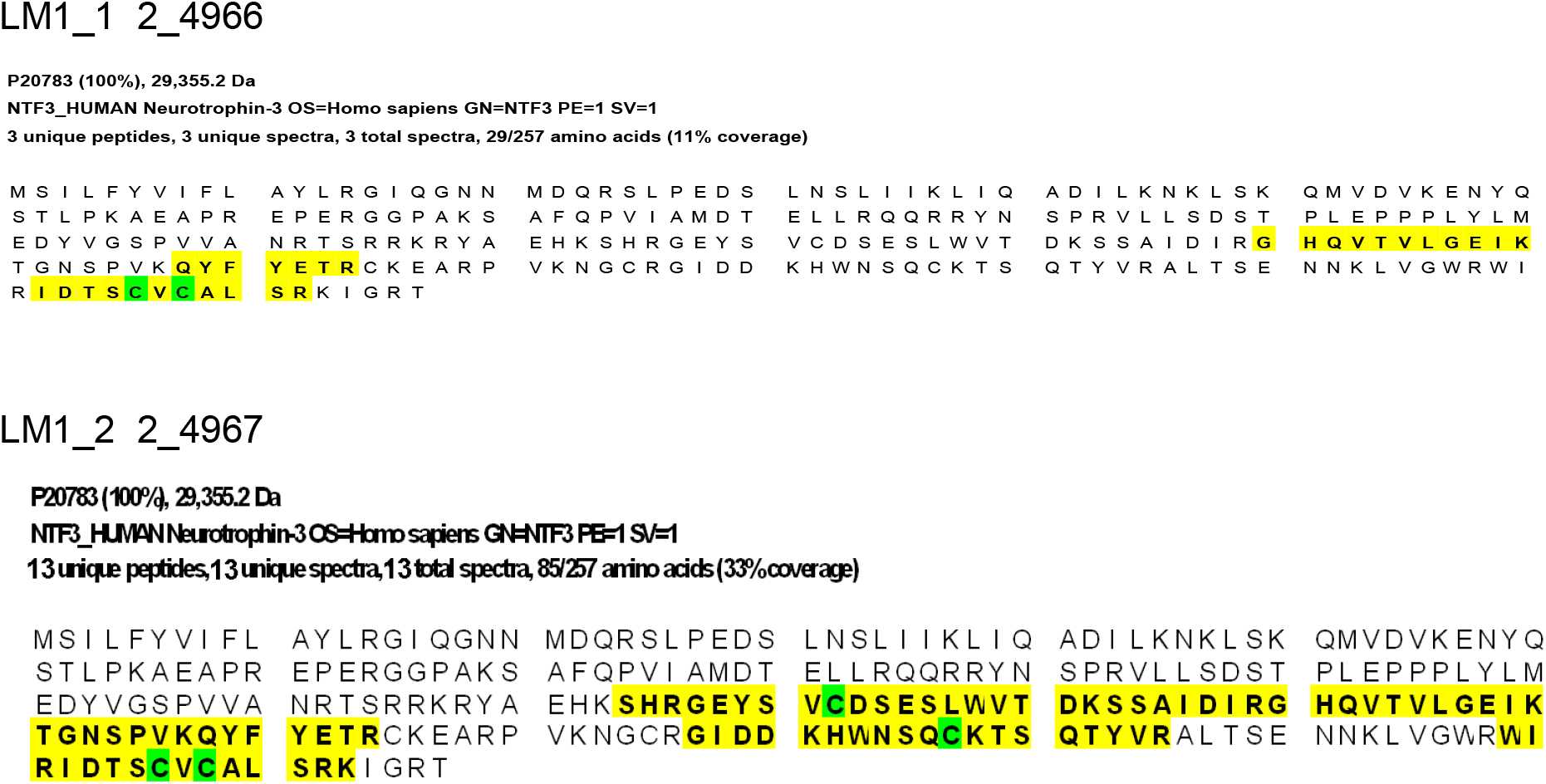
Proteomic analysis (LC/MS/MS sequencing) confirmed that the two bands were mature human neurotrophin-3 which begins YAEHK (seen just upstream of the first yellow region) and ends SRKIGRT (seen at the end of the yellow region). Coverage corresponds to the mature form of NT3 obtained after cleavage of human prepro-neurotrophin-3, isoform 2 (NP_002518.1).

## Notes

The authors declare that no conflict of interest exists.

